# Sourdough starter-inspired living materials grown from probiotic consortia with engineered division-of-labor

**DOI:** 10.1101/2025.10.09.681537

**Authors:** Shuchen Wang, Yiqi Xu, Yuewei Zhan, Martin Echavarria Galindo, Sherman Cheuk Yui Chau, Liang Cui, William M. Shaw, Wen Jun Xie, Hor Yue Tan, Chengyuan Lin, Xue Jiang, Zhaoxiang Bian, Peter Dedon, George M. Church, Timothy K. Lu, Tzu-Chieh Tang, Yong Lai

## Abstract

Genetic circuits enable precise control over cellular behavior. As circuits become more complex, implementing them within a single cell population becomes difficult. Tasks can instead be distributed in engineered consortia. However, selecting appropriate consortia and harnessing the natural traits of each species for such a division of labor remain unsolved challenges. Here, we report the creation of SINERGY, an engineered synthetic sourdough starter in which functional modules are distributed between the two constituent species. These modules, consisting of sensing, communication, and response, enable *in vitro* biosensing and *in vivo* drug delivery and modulate the gut ecosystem in an animal disease model. SINERGY not only preserves the safety and health-promoting properties of the natural components but also endows microbes with programmable functionalities, facilitating broad applications in biomedicine.

## Introduction

In synthetic biology, gene circuitry underlies the design and implementation of various functions in living cells. Over the last two decades, synthetic biologists have engineered microbes by deploying genetic circuits for biosensing (*1*, *2*), biocomputing (*3*), memory retention (*4*), reporting (*5*), and bioproduction (*6*). Genetic circuits, consisting of protein-coding genes and regulatory elements, are typically implemented in a single microbial population to perform an assigned function. Early synthetic genetic circuits consisted of simple on-off switches or basic logic gates; the pioneering achievements of the genetic toggle switch and the synthetic repressilator demonstrated the first programmable bistable switching and sustained rhythmic gene expression in living cells (*7*, *8*). These units have been combined and layered to construct complex circuits having sophisticated functions (*9*, *10*).

Nonetheless, implementing complex tasks within a single microbial population poses a significant challenge. As genetic circuits become more complex, there is a greater possibility of genetic interference and crosstalk within the cell, and resources may be limited (*11*). Coordinating the expression of numerous genes and regulatory elements while avoiding unintended consequences requires precise engineering and extensive optimization. Most importantly, the engineering capacity is limited by the nature of the microbial chassis, that is, the organism that harbors the genetic circuits (*12*). To overcome these limitations, complex tasks can be distributed among diverse microbial populations, so that each type of microbe performs some, but not all, of the desired genetic functions (*13*). Such a division of labor in engineered microbial consortia can reduce the burden and crosstalk of genetic circuitry placed on any one microbial type and simplify circuit optimization (*14*).

A stable symbiotic microbial consortium is essential for engineered task specialization, to manage resource competition and maintain long-term stability in a dynamic environment (*13*). In naturally evolved microbial communities, different subpopulations perform different tasks, modeling task distribution (*15*, *16*). Natural living materials comprising more than one species harness the capabilities of specialized cells in resource-limited conditions via a division of labor, ensuring survival and functioning (*17*, *18*). Lichen survives in harsh environments because its constituents, filamentous fungi and photosynthetic algae or cyanobacteria, exchange metabolites and nutrients (*19*). Similar processes occur in SCOBY (symbiotic community of acetic acid bacteria and yeasts), whose microbial metabolism produces kombucha tea (*20*). These stable natural consortia offer microbial chassis that can be used for a division of labor among engineered functions.

Engineered living materials (ELMs), consisting of microbes engineered to grow as living biofilms or to interface with non-living materials (*21–25*), respond to their environment and have multiple programmable functional properties (*26–29*). Whereas most ELMs consist of single strains (*30*), Gilbert et al., inspired by SCOBY-fermented kombucha beverages, co-cultured genetically engineered *Saccharomyces cerevisiae* (*S. cerevisiae*) with cellulose-producing *Komagataeibacter rhaeticus* bacteria, creating syn-SCOBY (*31*), a self-assembled stable living material useful for biosensing. McBee et al., isolated a dominant bacterial strain of *Pantoea agglomerans* (*P. agglomerans*) that stably coexists with a fungal mycelium (*32*). Engineered *P. agglomerans* was combined with the mycelium to produce sensing and responsive composites. However, in these examples only a single microorganism was genetically engineered. Harnessing the full potential of stable symbiotic microbes requires simultaneously engineering the multiple organisms that constitute these communities. This way, tasks can be distributed and the resulting living materials can acquire enhanced stability and multifunctionality.

Traditional fermented foods consist of stable microbial ecosystems that provide remarkable models for engineering macroscopic living materials having a division of labor (*33*, *34*). The safety of fermented foods has been established by centuries of use, and they have been found to have beneficial effects on human gut health, helping to maintain gut homeostasis and improving digestion (*33*). Sourdough bread, in particular, is recommended during the onset of inflammatory bowel disease (IBD) and irritable bowel syndrome (IBS) because individuals with sensitive digestive systems may tolerate it better than other foods (*35*, *36*).

Here, we present <u>S</u>ourdough-<u>IN</u>spired <u>E</u>ngineering of <u>R</u>econstructed <u>G</u>ut-beneficial bacteria and <u>Y</u>east consortia (SINERGY). We rebuilt a synthetic sourdough starter consisting of a stable community of coexisting probiotic strains of *S. cerevisiae* and *Lactiplantibacillus plantarum* (*L. plantarum*), distributed optimized functional modules in these co-existing microbes, and engineered an independent cross-kingdom communication system to coordinate their behavior. We then evaluated SINERGY in a mouse model of IBD: SINERGY detected a disease-associated biomarker in the gut and expressed diagnostic and therapeutic molecules *in situ*, demonstrating potential functions in diagnosis and intervention. We anticipate that the strategies and tools used to transform the synthetic sourdough starter into SINERGY will facilitate the construction of novel ELMs starting from microbial consortia.

## Results

### Reconstruction of a stable, reproducible synthetic sourdough starter

Fermenting sourdough is an ancient practice. In sourdough starter, a live culture made of flour and water, stable microbial consortia grow on starch and gluten. Microbial communities in 500 sourdough starters from four continents exhibit consistent patterns of species dominance and co-existence, though temperature, humidity, flour composition, and bacterial and yeast strains vary (*37*). Lactic acid bacteria and yeast, the dominant components of natural sourdough starters, shape community structure (*37–39*). To recreate a stable microbial consortium as the foundation for a division of labor, we used well-defined components to establish a synthetic sourdough starter and investigated the interactions among constituent microbes in this reproducible and controllable system (**Fig. 1A**).

**Fig. 1:**
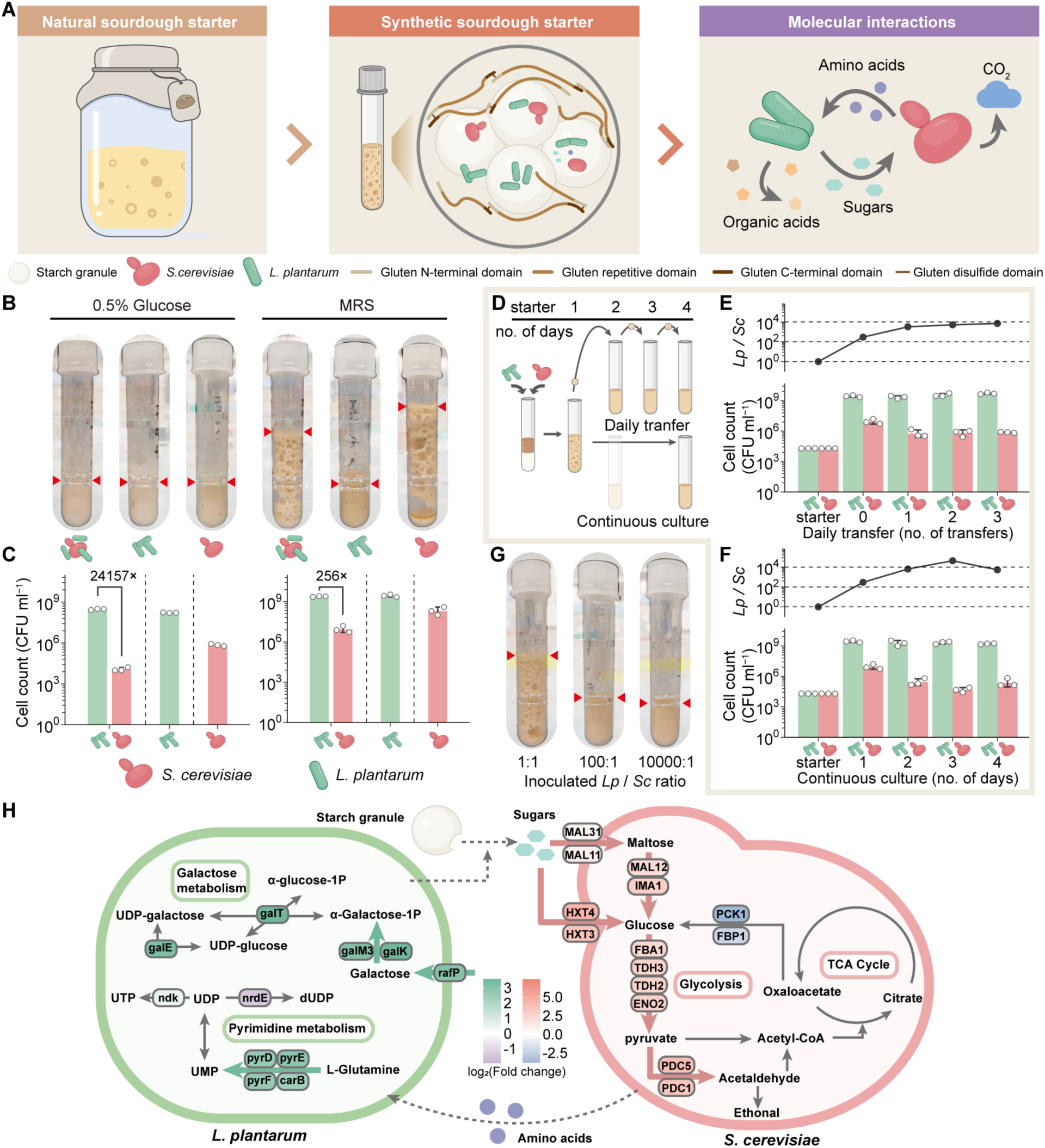
Reconstruction of sourdough-inspired synthetic starter. (**A**) Establishing and investigating a sourdough-inspired synthetic starter system under laboratory conditions. (**B** and **C**) Images (**B**) and cell counts (**C**) of synthetic starter grown from various liquid media (mean ± sd, n = 3 starters). The red triangular arrows indicate the fermentation height of the synthetic starter. (**D**) Daily transfer and continuous culture assay of synthetic sourdough starter. (**E** and **F**) Cell counts and ratio of *L. plantarum* WCFS1 to *S. cerevisiae* (*Lp* / *Sc*) over daily transfer (**E**) and days of continuous culture (**F**) (cell count represented by mean ± sd, ratio represented by mean value, n = 3 starters). (G) Images of the synthetic starter inoculated with different ratios of *L. plantarum* and *S. cerevisiae*. (H) Hypothetical interactions between *L. plantarum* and *S. cerevisiae* revealed by RNA-seq and metabolomics, by comparing co-culture with mono-culture conditions.

We reconstructed a sourdough-inspired starter by inoculating the commonly used laboratory strains *L. plantarum* WCFS1 or NC8 and *S. cerevisiae* BY4741 in synthetic “flour,” consisting of potato starch (90%) and gluten (10%). To support growth, we supplemented the synthetic starter with glucose as a basic carbon source and either YPD/YPS medium for cultivating yeast or MRS medium for cultivating *L. plantarum* (**Fig. 1, B and C,** and **fig. S1A**). When *S. cerevisiae* grew with or without *L. plantarum*, the synthetic starter with MRS medium exhibited the porous structure characteristic of natural sourdough fermentation and a marked dough rise (**Fig. 1B**). The porous structure and dough rise, primarily resulting from yeast fermentation and CO_2_ production, indicate the robust metabolic activity of yeast in a healthy fermentation state; this activity reduces gluten and FODMAPs (Fermentable Oligo-, Di-, Mono-saccharides And Polyols) (*40*). Although the initial inoculation ratio was 1:1, the quantity of *L. plantarum* was significantly higher than that of *S. cerevisiae* (256-fold) 48 hours after co-inoculation (**Fig. 1C**). In contrast, 0.5% glucose alone did not support the robust growth of both species, resulting in a disproportionately higher abundance (24157-fold) of *L. plantarum* compared to *S. cerevisiae*. With YPD or YPS as the culture medium, the typical porous fermentation phenotype was not observed (**fig. S1A**). Microbial quantification revealed the presence of *Bacillus* spp., likely originating from the gluten product (**fig. S1A**). Thus, MRS medium, as a liquid base with starch and gluten, supported the growth of a dominant microbial consortium, recapitulating the fermentation phenotype of natural sourdough starters.

To determine the stability of the co-existing microorganisms, we serially passaged 1:1 mixtures of *L. plantarum* WCFS1 or NC8 strains and *S. cerevisiae* for three 24-hour transfers or continuously cultured them for 72 hours and measured growth (**Fig. 1D**). Consistent with a previous study that used whole wheat and all-purpose flour (*37*), the symbiotic microbial consortium maintained a ratio of approximately 10,000:1 (*L. plantarum* to *S. cerevisiae*) after the second transfer and continuous culture (**Fig. 1, E and F,** and **fig. S1, B and C**). A commercial flour used as powder medium resulted in a similar texture and growth pattern (**fig. S1, D to F**). Given that the final cell count ratio of *L. plantarum* to *S. cerevisiae* was about 10,000:1, we investigated the effect of inoculation ratio on the synthetic starter with MRS medium. We observed the dough rise and porous structure when the synthetic starter was inoculated with *L. plantarum* and *S. cerevisiae* in a 1:1 ratio but not at ratios of 100:1 or 10,000:1 (**Fig. 1G** and **fig. S1G**). Scanning electron microscopy (SEM) visualization of the synthetic starter validated that these two microorganisms cohabitated on the starch granules (**fig. S1H**).

To investigate the interactions between *L. plantarum* and *S. cerevisiae*, we analyzed their transcriptome and metabolome under co-culture conditions, with monocultures as controls. Transcriptomic analysis revealed differential gene expression in both constituent strains (**fig. S2, A to C**). *S. cerevisiae* pathways associated with carbohydrate metabolism and biosynthesis of amino acids and proteins were upregulated during co-culturing with *L. plantarum*, compared to monocultures (**fig. S2C**). Specifically, *S. cerevisiae* genes involved in transporting maltose and glucose and their catabolism to pyruvate and acetaldehyde were significantly upregulated in co-culture (**Fig. 1H**). The apparent increased availability of simple sugars in the synthetic starter implies that *L. plantarum* may contribute to starch degradation, as seen with natural sourdough microbiota (*41*). Interestingly, *L. plantarum* genes related to pyrimidine and galactose metabolism were upregulated during co-culturing with *S. cerevisiae*, indicating enhanced uptake and utilization of L-glutamine and galactose (**Fig. 1H**). This upregulation was confirmed by targeted metabolomic analysis of culture supernatants (**fig. S3**). These observations are congruent with a previous study of co-cultured yeast-*Lactobacillus* in grape juice, where nitrogen overflow from the yeast created a niche for the symbiotic bacteria (*42*).

### Genetic engineering of resident bacteria for controllable output in SINERGY

Distributing diverse functions among the organisms co-existing in stable microbial communities not only reduces the metabolic burden and genetic crosstalk of complex genetic circuits for each type of cell but also allows full exploitation of the unique biological traits of each consortium member. In our synthetic starter, *L. plantarum* is an effective chassis for functional transformation into an actuator, because: 1) it is significantly more abundant than *S. cerevisiae* in this co-culture; and 2) as a natural inhabitant of the gastrointestinal tract, it can serve as a mucosal delivery vehicle (*43*). To engineer the synthetic starter, we first designed, characterized, and validated a range of biochemical interfaces, including modules for sensing and responding to external signals, in *L. plantarum* with inducible gene expression, logic processing, and output expression and secretion (**Fig. 2A**). Given that sourdough-inspired materials consist of 80-90% starch and that glycolysis is upregulated in co-cultured *S. cerevisiae*, we engineered *L. plantarum* to express α-amylase from *Lactobacillus amylovorus*: α-amylase catalyzes the hydrolysis of starch, which might change the macroscopic properties of the synthetic starter. To enhance amylase secretion and function, we tested the functionality of several signal peptides, short N-terminal amino acids that govern protein secretion. These peptides, sp2958 and spAmyL, significantly increased the secretion and activity of α-amylase, altering the texture and height of the synthetic starter (**Fig. 2, B and C**). Rheology measurement indicated less mechanical strength for the starter inoculated with the active amylase-secreting strain than for that inoculated with the wild-type strain (**Fig. 2D**), consistent with reports that amylase decreases the rheological properties of dough (*44*). As a control, we constructed a yeast strain that secretes α-amylase with the commonly used signal peptide MFα (α-mating factor), but it had weak enzyme activity and texture-changing efficacy (**fig. S4**), indicating the advantage of using engineered *L. plantarum* as the actuator in the synthetic starter.

**Fig. 2:**
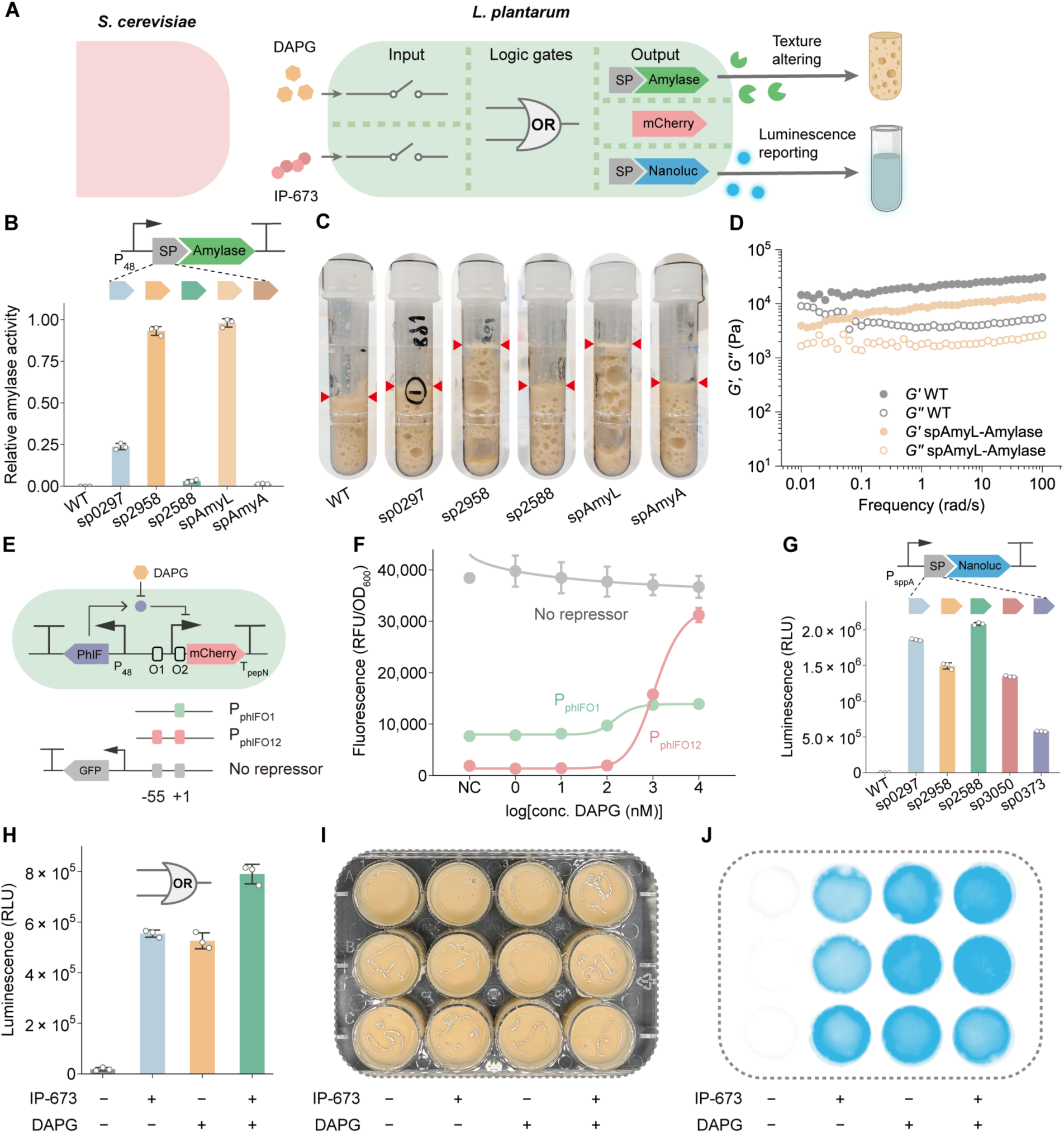
Design and characterization of the biochemical interfaces in *L. plantarum* as an actuator in SINERGY. (**A**) Schematic of the engineered *L. plantarum* in sensing, processing logic, and executing responses in SINERGY. SP: signal peptide. (**B** and **C**) Secretion capacity (**B**) and starter height (**C**) of *L. plantarum* strains that secreted amylase in response to 5 different signal peptides (mean ± sd, n = 3 independent cultures). The red triangular arrows indicate the fermentation height of the synthetic starter. (**D**) Frequency-dependent rheology properties of synthetic sourdough starter grown from WT or amylase-secreting *L. plantarum* led by spAmyL. (**E** and **F**) Design (**E**) and characterization (**F**) of DAPG-responsive *L. plantarum* strain (mean ± sd, n = 3 independent cultures). PhlF operator sites were inserted in regions O1 and O2 of the P_11_ promoter. NC, negative control. (**G**) Screening signal peptides for secreting NanoLuc luciferase (mean ± sd, n = 3 independent cultures). (**H**) An OR-gate construct in *L. plantarum* for sensing IP-673 or DAPG in liquid culture (mean ± sd, n = 3 independent cultures). (**I** and **J**) Bright field (**I**) and luminescence (**J**) images of OR-gate *L. plantarum* strain in SINERGY cultured in 12-well plates.

To enhance the response of *L. plantarum* to external molecules, we designed a system inducible by 2,4-diacetylphloroglucinol (DAPG). The PhlF repressor was expressed from the constitutive P_48_ promoter, and pairs of PhlF operator sites were inserted in the strong promoter P_11_ at O1 and O2 (**Fig. 2E** and **fig. S5A**). We assessed the performance of the DAPG-inducible promoter by using it to drive the expression of the fluorescent reporter mCherry. Without DAPG, the PhlF protein bound to the designed promoters, repressing mCherry expression (**Fig. 2F**). DAPG elicited 1.8- and 16.5-maximal fold increases in fluorescence output level for the two designs, respectively, with a sensitivity of 0.23 µM for the P_phlFO12_ design.

To further quantify the secretion and stimulus-responsive capabilities of the synthetic starter, we engineered *L. plantarum* to secrete NanoLuc luciferase. In this strain, NanoLuc expression is driven by a sakacin P peptide IP-673-inducible system (**fig. S5B**) under the control of the inducible promoter P_sppA_ (*45*). We then assessed the effects of five signal peptides on NanoLuc secretion efficiency (**Fig. 2G**); of these, sp2588 resulted in the highest secretion efficiency. To evaluate the capacity of *L. plantarum* to perform basic logic processing, we designed two OR gates by integrating the DAPG- and peptide-inducible systems (**fig. S5C**). The promoter architecture in Design 2 effectively implemented OR gate functionality (**fig. S5D**). In both liquid culture (**Fig. 2H**) and synthetic starter (**Fig. 2, I and J**), peptide and small molecule inducers activated NanoLuc expression, whereas without induction, leaky expression remained minimal. Thus, we constructed, characterized, and optimized the key functional modules of *L. plantarum* as an actuator.

### Genetic engineering of eukaryotic cells for sensing and communication in SINERGY

Communication between eukaryotic and prokaryotic cells is essential to create layered circuits for the systems-level applications of materials (**Fig. 3A**). As the eukaryotic component in our system, *S. cerevisiae* has several key attributes that make it a good choice for transformation into the chassis of a living biosensor: 1) it accommodates eukaryotic sensing modalities, such as G-protein-coupled receptors (GPCRs), expanding the range of detectable signals; 2) it thrives in harsh environments; and 3) it can be produced cheaply in ‘active dry’ form and remains viable for extended periods.

**Fig. 3:**
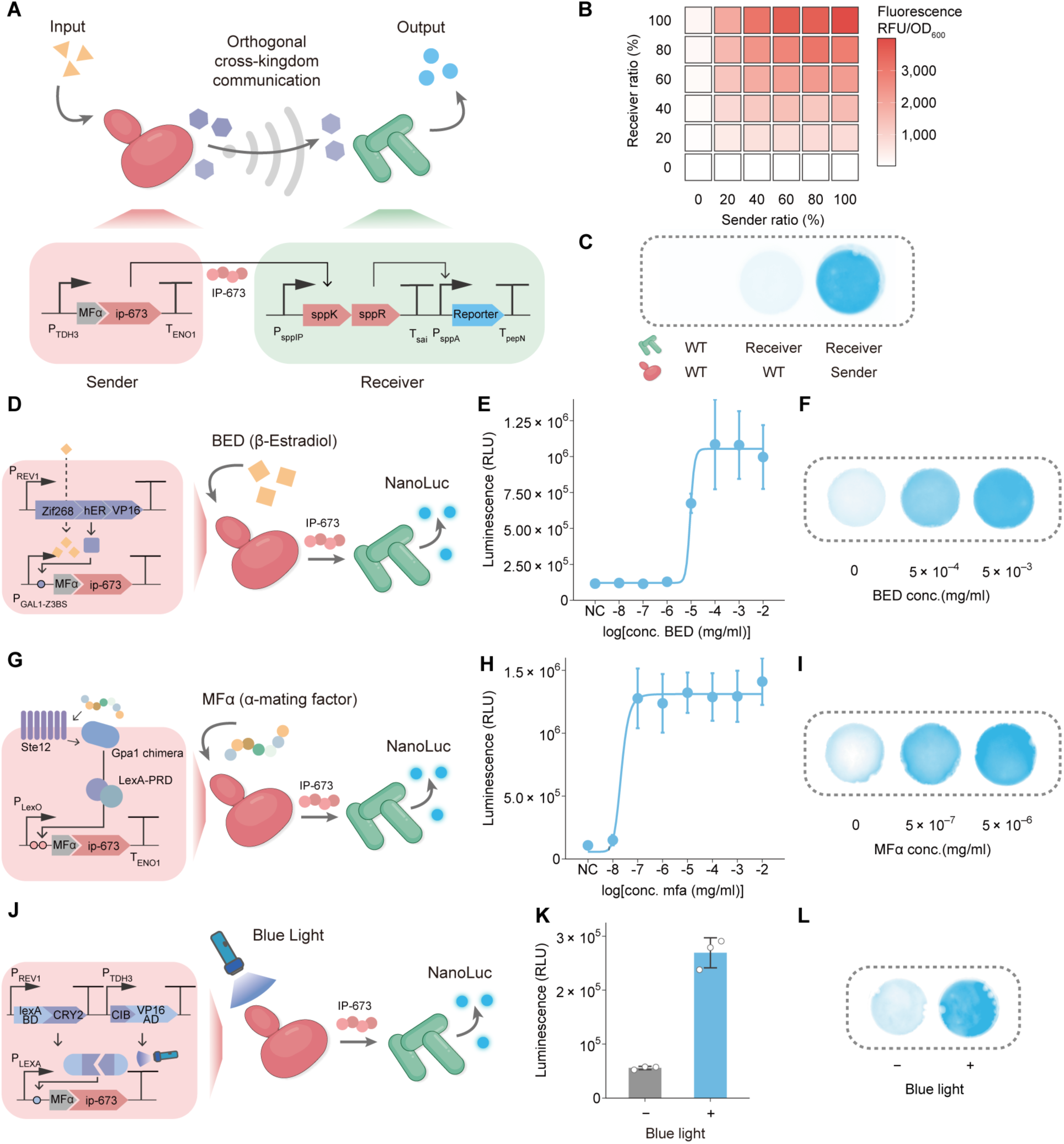
An orthogonal cross-kingdom communication system bridges an *S. cerevisiae* sensor and an *L. plantarum* actuator. (**A**) Schematic of the engineered division of labor in *S. cerevisiae* (the sender, in pink) and *L. plantarum* (the actuator, in light green). (**B** and **C**) Characterizing the IP-673-mediated division of labor in liquid medium (RFU, relative fluorescence units) (**B**) and synthetic starter (**C**). (**D**-**L**) The response of engineered microbial consortia to various signals in liquid medium and synthetic starter: BED (**D**-**F**), MFα (**G**-**I**), and blue light (**J**-**L**). NC, negative control. (B is represented as mean value; E, H, and K are represented as mean ± sd, n = 3 independent cultures).

The sakacin P peptide IP-673-inducible system, having a low background and a high inducing level, has been developed in *L. plantarum*. To create SINERGY, we re-purposed the inducing peptide IP-673 as a secreted output of *S. cerevisiae* guided by the MFα signal peptide, which triggers downstream responses in *L. plantarum*. When both strains were inoculated in MRS medium, the fluorescence output increased as the relative proportion of either the yeast sender strain or the *L. plantarum* receiver strain in the population rose relative to its respective wild-type strain (**Fig. 3B**). This result showed that engineered *S. cerevisiae* could express and secrete peptides at sufficient concentrations to modulate gene expression in *L. plantarum*, thereby mediating cross-kingdom communication. To validate the function of the communication system in synthetic starter, we co-cultured the yeast sender strain and a NanoLuc-secreting receiver strain of *L. plantarum* for 24 hours in 12-well plates. The luminescence produced by SINERGY was higher than in control groups, demonstrating that communication had been established between the eukaryotic and prokaryotic cells (**Fig. 3C**).

We engineered yeast biosensor strains to secrete peptide IP-673 as the cell-to-cell communication signal for detecting: 1) a transcriptional factor-based sensor in response to β-estradiol (BED) (**Fig. 3D**); 2) a GPCR-based sensor in response to MFα (**Fig. 3G**); and 3) a blue-light sensor (**Fig. 3J**). Upon sensing these stimuli, *S. cerevisiae* produced and secreted IP-673. This peptide was sensed by *L. plantarum*, which then quantifiably expressed NanoLuc. We first characterized the dose-response ratio of these systems by co-culturing yeast and *L. plantarum* in MRS broth. The consortium responded to corresponding stimuli in liquid cultures by emitting either luminescence or fluorescence (**Fig. 3, E, H, and K** and **fig. S6)**. Luminescence imaging indicated the division of labor between consortium members detecting external signals (*S. cerevisiae*) and responding with NanoLuc production (*L. plantarum*) in SINERGY (**Fig. 3, F, I, and L**).

### High-throughput forward genetics in *L. plantarum* to identify bacterial fitness communities

Fermented foods serve as a significant dietary reservoir of microbes, potentially benefitting human health (*46*, *47*). Yet, the direct delivery of probiotics may be constrained by the acidic stomach environment, bile acid produced in the gut, and other difficulties in engraftment (*48*). To improve *in vivo* delivery efficiency by accelerating probiotic adaptation to the gut environment, we employed a forward genetics approach.

We first evaluated the survival of probiotics, delivered by the synthetic starter, in the stool (**Fig. 4A**) and the murine digestive tract (small intestine, cecum, and large intestine) (**fig. S7, A and B**). At 24 hours post-gavage, the abundance of *L. plantarum* and *S. cerevisiae* was significantly reduced compared to the 6-hour timepoint, with *L. plantarum* greatly outnumbering yeast. Interestingly, at 48 hours, the number of *L. plantarum* cells was higher than that in the PBS control group, suggesting that the synthetic starter had promoted bacterial persistence in the gut (**Fig. 4A**). The fluorescence emitted in the murine gut by the engineered organisms was also evaluated. Confocal imaging of the colon 3 hours post-administration showed localized mCherry-expressing *L. plantarum* and GFP-expressing *S. cerevisiae* (**Fig. 4B** and **fig. S7, C and D**).

**Fig. 4:**
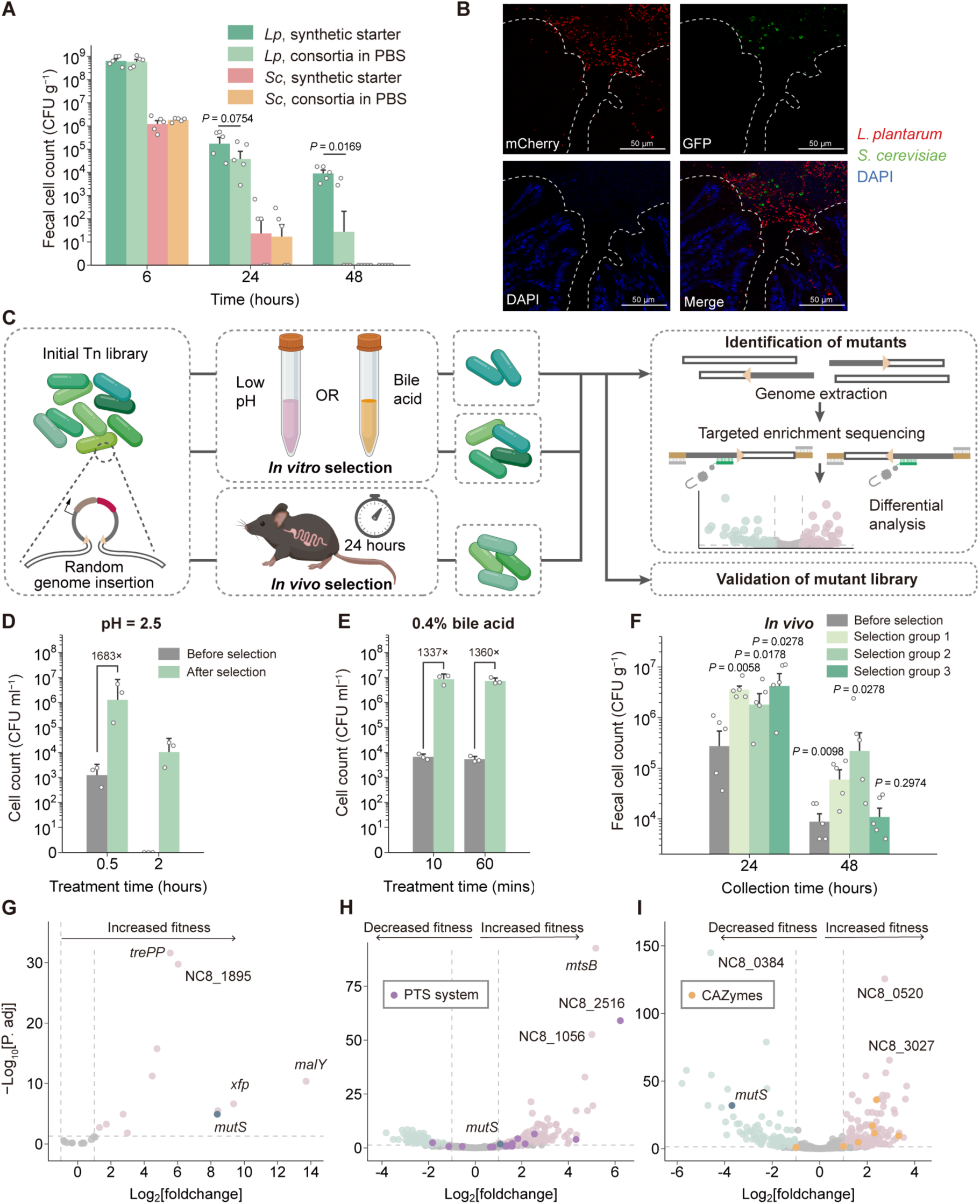
Transposon insertion sequencing enables high-throughput forward genetic screen to select bacterial fitness community. (**A**) Fecal cell counts of *L. plantarum* and *S. cerevisiae* delivered to mice via synthetic starter or PBS (mean ± SEM; n = 5 mice; statistical significance was determined by Mann–Whitney test). Mouse fecal samples were collected at 6, 24, and 48 hours post-gavage. (**B**) Confocal images of mouse colon sections 3 hours post-gavage with synthetic starter. *L. plantarum* expresses mCherry, *S. cerevisiae* expresses GFP, and DAPI stains eukaryotic nuclei. (**C**) Schematic of large-scale transposon mutagenesis, selection, and analysis. (**D**-**F**) Validation of the selected *L. plantarum* mutant library. Cell counts of *L. plantarum* mutant library after selection compared with the original library in *in vitro* acidic environments (**D**) (mean ± sd, n = 3 independent experiments) and in bile acid salt (**E**) (mean ± sd, n = 3 independent experiments). Fecal cell counts of *L. plantarum* mutant library delivered via synthetic starter after selection in the intestinal tracts of mice compared with the original library (**F**) (mean ± SEM; n =5 mice; statistical significance was determined by Mann–Whitney test compared with group before selection of the same time points). (**G**-**I**) Identification of the selected *L. plantarum* mutants. Volcano plot of insertions from evolution groups versus original library selected in acidic environments (**G**), bile acid salt (**H**), gut residence time (**I**). Increased-fitness insertions and decreased-fitness insertions are labeled as purple and green dots, respectively. Grey dots represent non-significant insertions. For each selection, experiments were carried out in triplicate.

To identify mutants with greater fitness than wild-type *L. plantarum*, we constructed an IS1223 transposase-based transposon mutagenesis system (*49*, *50*) and built a large-scale mutant library (**fig. S8A**). We used targeted enrichment next-generation sequencing to accurately track insertion sites, confirming the coverage, within the library, of 1279 genomic open reading frames (ORFs, 42% of *L. plantarum* NC8 annotated ORFs) and 506 intergenic regions (**fig. S8B**). To assess the feasibility of the forward genetic approach in *L. plantarum*, we selected, by three rounds of *in vitro* treatments, mutant populations that adapted to gastric acidity (low pH) and intestinal bile acids (**Fig. 4C** and **fig. S8, C and D**). The fitness after selection was validated, and the genotypes were analyzed by sequencing. Under acidic conditions (pH=2.5) mutants selected by this approach were three orders of magnitude more abundant than in the unselected population, with enriched mutations in genes such as *malY* (a putative aminotransferase), *xfp* (xylulose-5-phosphate/fructose-6-phosphate phosphoketolase), *trePP* (a glycosyl hydrolase) and NC8_1895, encoding a hypothetical protein (**Fig. 4, D and G**). Similarly, mutants selected under 0.4% bile acid stress demonstrated enhanced tolerance: mutations occurred in cation permease *mtsB*, which is an ATP-binding cassette (ABC) transporter, NC8_2516, predicted to be associated with the phosphotransferase system (PTS), and NC8_1056, encoding a hypothetical protein (**Fig. 4, E and H**). We next applied this high-throughput forward genetics strategy in an *in vivo* context, selecting sub-populations that resided for a longer time in the mouse gut (**Fig. 4, F and I**). Following three rounds of selection in the intestinal tracts of three independent groups of mice, *L. plantarum* strains recovered from feces exhibited adaptive mutations enriched in carbohydrate-active enzymes (CAZymes) and several genes encoding hypothetical proteins (i.e., NC8_0520 and NC8_3027), suggestive of improved fitness. Interestingly, the dysfunction of the *mutS* gene, which recognizes mismatches and contributes to DNA replication fidelity, led to higher fitness in high bile acid and low pH conditions, while decreasing the fitness for mouse engraftment, illustrating the greater importance of replication fidelity *in vivo* than *in vitro*. Those *in vivo*-selected mutant populations pointed to the rational design of microbial consortia optimized for gut delivery.

### Engineering division of labor in SINERGY for sensing and reporting gut inflammation

As fermented foods can serve as vehicles for delivering symbiotic microbes to the gut, we set out to engineer SINERGY to dynamically sense and respond to gut inflammation signals in a self-tunable and division-of-labor manner (**Fig. 5A**).

**Fig. 5:**
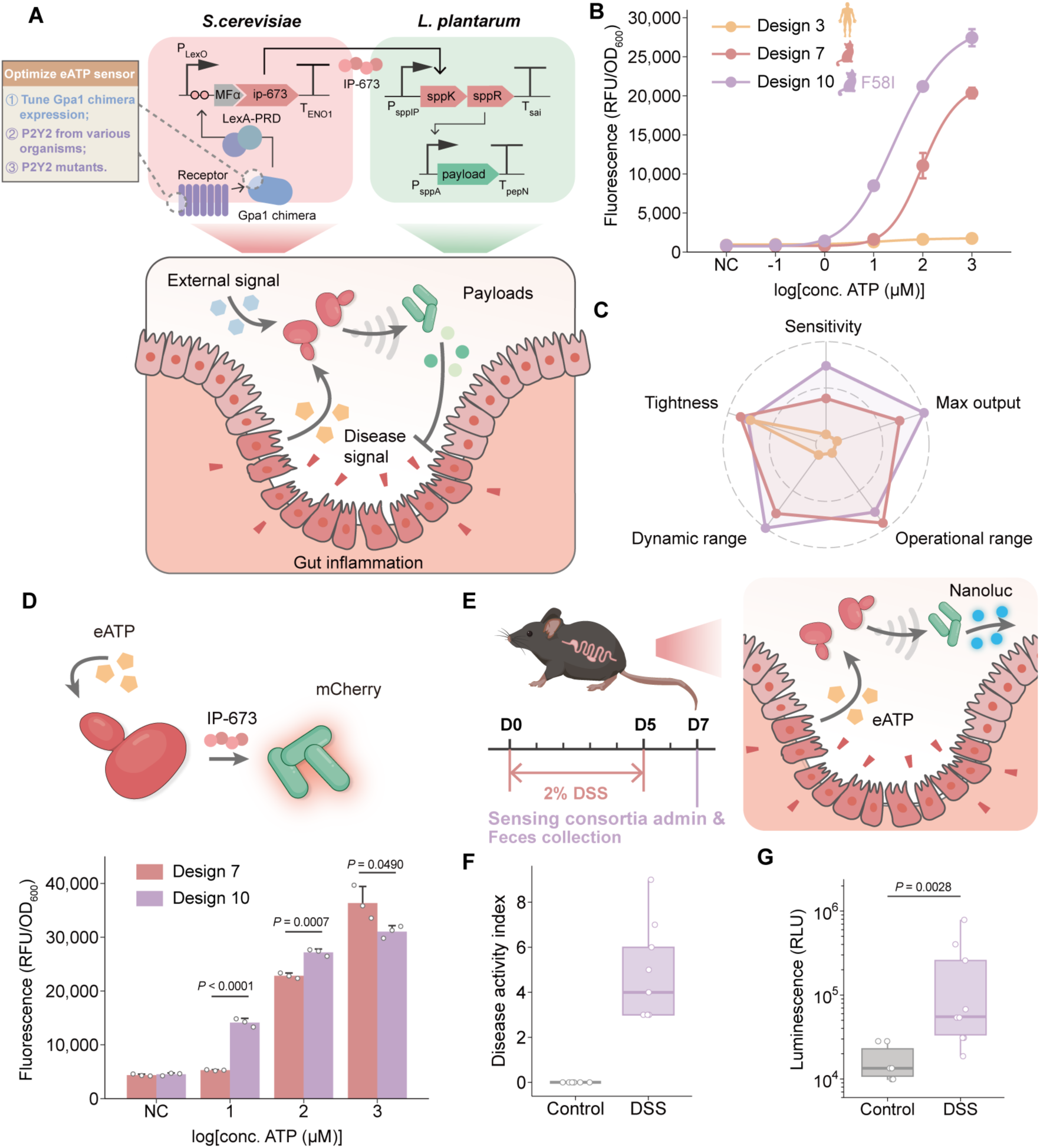
Engineering SINERGY to sense and respond to gut inflammation. (**A**) Schematic of SINERGY for sensing and responding to gut inflammation and strategies for engineering yeast GPCR biosensors to detect eATP (extracellular ATP). (**B**) Dose-response curve of representative eATP sensor designs (mean ± sd, n = 2 or 3 independent cultures). For Designs 3 and 7, the DNA sequence of the P2Y2 receptor was derived from human and cat, respectively. Design 10 has an F58I point mutation in the cat P2Y2 receptor. (**C**) Characterizing sensing properties for representative eATP sensor designs. (**D**) Characterizing eATP-sensing consortia with *L. plantarum* expressing mCherry as output in MRS broth. Fluorescence was measured after 6 hours of ATP induction (mean ± sd, n = 3 independent cultures; statistical significance was determined by unpaired two-sided t-test). (**E**) Mouse experiments for verification of inflammation-sensing consortia. (**F** and **G**) Disease activity index (**F**) and relative luminescence units (**G**) of fecal samples harvested from DSS and control groups (n = 6 mice for Control group, n = 9 mice for DSS group, statistical significance was determined by Mann-Whitney test).

Extracellular adenosine triphosphate (eATP) generated by both the commensal microbiota and host cells triggers purinergic signaling, which enhances intestinal inflammation and contributes to pathology (*51*, *52*). By incorporating the GPCR sensing capacity of *S. cerevisiae*, we engineered an eATP sensor expressing the P2Y2 receptors to detect gut inflammation. Scott et al. developed an eATP reporter strain based on the *S. cerevisiae* CB008 strain, but its construction required multiple steps of genome manipulation (*53*). To construct a modular eATP sensing-sender yeast strain more easily, we started with yWS677, a model *S. cerevisiae* strain refactored for GPCR signaling (*54*). This BY4741-derived yWS677 strain, expressing the human P2Y2 receptor and signaling either by the original *STE12-pFUS* pathway (Design 1) or by the *LexA-PRD* transcription factor targeting a synthetic promoter (Design 2), did not respond to eATP (**fig. S9A**). We then designed and optimized eATP biosensors through different strategies (**Fig. 5A** and **fig. S9A**) and evaluated their sensitivity, maximum output, operational range, dynamic range, and tightness. We used a weaker promoter (P_ALD6_) to tune down Gpa1 chimera expression (Design 3) and observed a slight response to ATP. We then tested P2Y2 from other organisms (mouse, olive baboon, green monkey, cat; Designs 4-7) with the same construct; intriguingly, cat P2Y2 exhibited the highest response (Design 7). To maximize sensor sensitivity, we generated three mutations on cat P2Y2 (Designs 8-10) that increase the response of human receptors (*53*). Mutation F58I on cat P2Y2 generated higher sensitivity, dynamic range, and output level in response to ATP stimuli than mutation Q165H or N116S (**Fig. 5, B and C** and **fig. S9, A and B**). Finally, we screened promoters for expressing Gpa1 chimera for cat P2Y2 F58I. The design using P_ALD6_ to control the expression of Gpa1 chimera had the most sensitivity and satisfied responsiveness (Design 10; **fig. S9C**).

Consortia consisting of swF221 (NanoLuc-secreting *L. plantarum* receiver strain) and a yeast sensor strain (either Design 7 [wild-type cat P2Y2] or Design 10 [cat P2Y2 F58I]) responded well to pathological eATP concentrations after 6 hours of induction; the latter consortium exhibited greater sensitivity (**Fig. 5D**). We then tested the eATP-sensing consortia *in vivo* in dextran sulfate sodium (DSS)-treated and healthy control mice (**Fig. 5E**). Luciferase levels in the DSS-treated group were significantly elevated with the swF221 *L. plantarum* strain and Design 10 *S. cerevisiae* strain compared with healthy controls, corresponding with the raised Disease Activity Index (DAI) (**Fig. 5, F and G**).

### Therapeutic SINERGY alleviates DSS-induced colitis

Anti-inflammatory cytokines and serine protease inhibitors help prevent overwhelming gut inflammation, strengthen epithelial barrier functions, and restore gut homeostasis (*55*, *56*). As SINERGY could sense eATP in the murine gut, we explored whether the stably co-existing *L. plantarum* could function as an actuator for the *in situ* secretion and delivery of therapeutic payloads (**Fig. 6A**). To assay for the recovery of gut homeostasis as the output, we chose 3 therapeutic payloads, each led by four signal peptide candidates, to be secreted by *L. plantarum*: the murine anti-inflammatory cytokine IL-10 (mIL-10), the human serine protease inhibitor Elafin (hElafin), and the murine secretory leukocyte peptidase inhibitor (mSLPI). The signal peptides for mIL-10 and hElafin had outstanding secretion capacity (**Fig. 6B**). Secretion was implemented in the *L. plantarum*-adapted community (*in vivo* selection group 1) having the longest *in vivo* residence time in the murine gut (**Fig. 4F**).

**Fig. 6:**
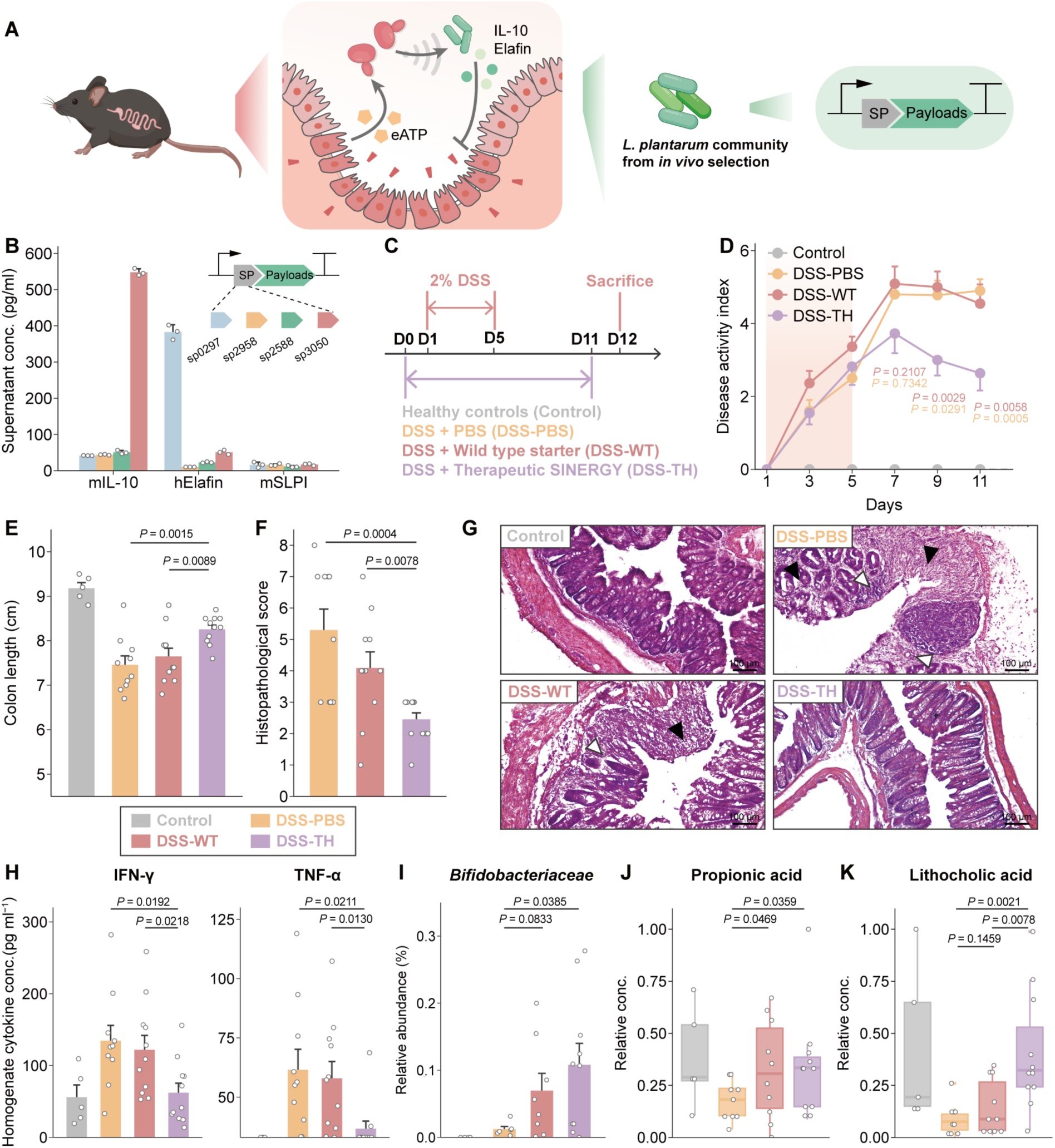
Alleviation of DSS-induced colitis by therapeutic SINERGY consisting of engineered consortia. (**A**) Schematic of therapeutic consortia for alleviating IBD, in which an *L. plantarum* community selected by *in vivo* screening was engineered for therapeutic payload secretion. (**B**) Quantification of the secretion capacities for mIL-10, hElafin, and mSLPI by ELISA (mean ± sd, n = 3 independent cultures). (**C**) Schematic of experimental design and grouping. (**D**) Disease activity index of mice in each group (mean ± SEM, statistical significance was determined by two-way ANOVA. P value representing comparison of DSS-TH group versus DSS-PBS or DSS-WT group, n = 5 mice for Control group, n = 10 for DSS-PBS group, n = 11 for DSS-WT and DSS-TH group). (**E** and **F**) Colon length (**E**) and histopathological scoring (**F**) for each group of mice (mean ± SEM, statistical significance was determined by unpaired two-sided t-test, n = 5 mice for Control group, n = 10 for DSS-PBS group, n = 11 for DSS-WT and DSS-TH group). (**G**) Representative images of distal colon sections from each group with hematoxylin and eosin staining. Black arrow: epithelial changes, including disappearance or damage of glands and goblet cells; white arrow: lymphocytic infiltration. (**H**) Representative inflammatory cytokine levels of colon homogenate (mean ± SEM, statistical significance was determined by unpaired two-sided t-test, n = 5 samples for Control group, n = 10 for DSS-PBS group, n = 11 for DSS-WT and DSS-TH group). (**I**) Relative *Bifidobacteriaceae* abundance in feces (mean ± SEM, statistical significance was determined by unpaired two-sided t-test, n = 5 samples for Control group, n = 7 for DSS-PBS group, n = 8 for DSS-WT and n = 10 DSS-TH group). (**J** and **K**) Relative concentration of propionic acid (**J**) and lithocholic acid (**K**) of colon content (statistical significance was determined by unpaired two-sided t-test, n = 5 samples for Control group, n = 9 for DSS-PBS group, n = 10 for DSS-WT and n = 11 DSS-TH group).

We evaluated the therapeutic capacity of engineered SINERGY consortia, consisting of the eATP sensor *S. cerevisiae* strain (Design 10) and the *L. plantarum* strain secreting sp3050-mIL-10 and sp0297-hElafin in a DSS-induced model of colitis (**Fig. 6A**). We administered daily either therapeutic SINERGY (DSS-TH [therapeutic] group, n = 11 mice), PBS (DSS-PBS group, n = 10 mice), or synthetic starter with wild-type yeast and *L. plantarum* (DSS-WT [wild-type] group, n = 11 mice). A healthy control group was also included (Control group, n = 5 mice) (**Fig. 6C**). The DAI and colon length of the DSS-TH group exhibited significant improvement (**Fig. 6, D and E**). Intestinal bleeding in DSS-induced mice was most severe on the second day following the cessation of chemical induction (DSS-PBS and DSS-WT groups), whereas in the DSS-TH group, bleeding was significantly reduced (**fig. S10A**). The DSS-TH group lost less weight than the DSS-WT group, but no significant difference was observed between the DSS-TH group and the DSS-PBS group (**fig. S10B**). *L. plantarum* supplementation in the early stages of the intervention may have suppressed weight gain (*57–60*). Histological analysis of the distal colon section showed improved colorectal integrity in the DSS-TH group compared to the DSS-PBS and DSS-WT groups, for which epithelial damage and inflammatory infiltration were observed (**Fig. 6, F and G**). Given the immunomodulatory role of IL-10, we measured the levels of inflammation-associated cytokines in intestinal cells. Lower amounts of proinflammatory cytokines (IFN-γ, TNF-α, IL-6, IL-17A, and MCP-1) were produced by the intestinal cells of the DSS-TH group than by those of the DSS-PBS or DSS-WT groups, while IL-10 levels did not differ significantly (**Fig. 6H** and **fig. S10C**).

To assess the effect of therapeutic SINERGY on repopulating the gut microbiota and restoring its function, we analyzed the microbiota composition and the bacterial-derived metabolites in the mouse intestine. The relative abundance of *Bifidobacteriaceae* was significantly higher in the DSS-TH groups compared to either the diseased or the healthy control groups (**Fig. 6I**). The short-chain fatty acid (SCFA) propionic acid was significantly elevated in the colon of the DSS-TH group, suggesting that the synthetic starter supported the growth of probiotic bacteria and the production of beneficial metabolites (**Fig. 6J**). Although the abundance of *Lactobacillaceae* did not change significantly, *Lachnospiraceae*, which are associated with SCFA production, recovered in the DSS-TH group (**fig. S10, D and E**). The significant increase in several secondary bile acids (lithocholic acid, ursodeoxycholic acid, deoxycholic acid) and the decrease in primary bile acid (glycocholic acid) in the DSS-TH group provided additional evidence of restored microbial functionality (**Fig. 6K** and **fig. S10F**). More indole-3-propionic acid (IPA), which mitigates intestinal inflammation through the aryl hydrocarbon receptor (AHR), was also produced in the DSS-TH group compared to DSS-WT group (**fig. S10G**).

In summary, our engineered therapeutic SINERGY improved multiple disease indicators in a murine IBD model. SINERGY enabled the precise sensing of an IBD-associated biomarker and the *in situ* delivery of therapeutic payloads to enhance intestinal barrier integrity and modulate the immune response, thereby reducing inflammation and restoring gut homeostasis. SINERGY also promoted the recovery of beneficial functions of the native gut microbiota, which collectively contributed to its therapeutic efficacy (**fig. S10H**).

## Discussion

Centuries of human consumption and cultural transmission have shown the safety and health benefits of traditional fermented foods. The fermentation of sourdough starter reduces antinutritional factors that may induce inflammatory or immunogenic responses, enhances probiotic growth, and increases the availability of beneficial bioactive compounds (*40*). By reprogramming traditional fermented foods, we can both preserve their intrinsic health-promoting properties and endow symbiotic microbes with new functionalities, enabling applications ranging from *in vitro* biosensing to *in vivo* drug delivery and the modulation of gut ecosystems.

Here, we demonstrate that SINERGY, a synthetic sourdough starter grown from engineered probiotic consortia, can acquire complex sensing and response capabilities. By co-culturing microbes typically found in sourdough starters, we established an orthogonal communication system bridging the sensing circuits in the eukaryote *S. cerevisiae* with the response and secretion circuits in the prokaryote *L. plantarum*. This integration enables each microbe to perform specialized functions while contributing to a unified consortium with coordinated operation in a complex environment.

Genetic circuits constructed with a single microbial chassis have limited capacity for incorporating sophisticated functions or precise regulatory control. Distributing distinct genetic modules across multiple host strains reduces the circuit complexity within each chassis. By harnessing the microbial consortia naturally present in traditional fermented foods, our study demonstrated two fundamental design principles for engineering a division of labor.

The first is functional specialization: each constituent microbe leverages its unique, inherent, and complementary capabilities. The second is consortium stability: ELMs perform consistently in dynamic environments, such as the mammalian gut. Modularity in genetic circuit design simplifies construction and validation, enabling the efficient replacement of biosensing and payload components as well as tailored responses to distinct environmental signals.

To assess the functionality of SINERGY in the gut, we designed, built, and optimized GPCR-based biosensors in *S. cerevisiae* and built an actuator on an adaptively evolved *L. plantarum* community for payload production and delivery. In a murine model of IBD, SINERGY sensed an IBD-specific biomarker, triggered the secretion of multiple therapeutic molecules, modulated immune responses, and enhanced intestinal epithelial barrier function. This coordinated response mitigated excessive gut inflammation and restored gut homeostasis, thereby facilitating the recovery of the beneficial functions of the native gut microbiota. In the future, SINERGY may be encapsulated and developed to relieve the symptoms, or even causes, of inflammatory intestinal disease, offering advantages over conventional therapeutic strategies aimed at a single target. Therefore, we anticipate that SINERGY will facilitate the development of a broad range of applications in biology and medicine.

The ability to respond to supplemented therapeutic molecules, as well as sensing pathological signals, would be a valuable property as it would enable the external control of SINERGY, thereby enhancing its safety. As supplemented niacin inhibits pro-inflammatory genes in macrophages and decreases epithelial cell apoptosis in ulcerative colitis (*61*), we engineered another *S. cerevisiae* strain and incorporated it into a consortium for sensing niacin. The consortium, however, exhibited only a 3.4-fold output after 6 hours of induction by 1 mg/ml niacin, indicating insufficient sensitivity for therapeutic relevance (**fig. S9, D and E**). Consequently, we did not proceed with the *in vivo* validation of this construction. Nevertheless, SINERGY may eventually be developed to respond to niacin or other activators, and future studies equipping SINERGY with the ability to sense a broader spectrum of external signals may improve its responsiveness and the autonomous regulation of therapeutic payloads production.

## Materials and methods

### Strains, culture conditions and transformation

*Escherichia coli (E. coli)* DH5α, used to propagate and construct all plasmids unless specified, was grown in LB medium at 37°C with appropriate antibiotics (carbenicillin 100 µg ml^−1^, erythromycin 200 µg ml^−1^, chloramphenicol 34 µg ml^−1^, and kanamycin 50 µg ml^−1^). *L. plantarum* WCFS1 and NC8 strains were grown in MRS medium at 30°C statically, with appropriate antibiotics (erythromycin 10 µg ml^−1^, chloramphenicol 10 µg ml^−1^, tetracycline 40 µg ml^−1^). *S. cerevisiae* BY4741 MATa was grown in YPD medium or selective YNB+N-U medium with shaking at 30°C. Yeast GPCR-based sensor strains were constructed based on strain yWS677(*54*). YPD and YPS medium were prepared with 10 g l^−1^ yeast extract, 20 g l^−1^ peptone, and 20 g l^−1^ glucose or sucrose. YNB-U+N medium was prepared with 1.4 g l^−1^ yeast synthetic dropout medium, 6.8 g l^−1^ yeast nitrogen base without uracil, and 20 g l^−1^ glucose.

Plasmids were introduced into *L. plantarum* following the previously reported protocol with adjustments (*62*). An overnight culture was used to inoculate 5 ml of SGMRS (MRS supplemented with 0.75 M sorbitol and 1% glycine) at a concentration of 2% (v/v). The culture was incubated at 30°C until it reached an OD_600_ between 0.3 and 0.5. The cells were then collected by centrifugation at 4°C, washed twice with ice-cold SM buffer (952 mM sucrose, 3.5 mM MgCl_2_), and resuspended in 90 μl of the same buffer. Subsequently, 500 ng of plasmid DNA was introduced to the cell suspension. After mixing, the sample was kept on ice for 10 minutes, transferred to a 0.2-cm cuvette (Bio-Rad), and electroporated using a Gene Pulser (Bio-Rad) under conditions of 2000 V, 25 μF, and 400 Ω. Following electroporation, 1 ml of MRS broth was added to the cuvette, and the cells were recovered at 30°C for 3 hours. Finally, the cells were plated on MRS agar plates with appropriate antibiotics.

For *S. cerevisiae* transformation, the LiAc/SS carrier DNA/PEG method was used (*63*). 1 ml overnight culture of *S. cerevisiae* were harvested by centrifugation, washed two times with 1 mL of sterile 0.1 M lithium acetate (LiAc), and finally resuspended in 0.1 M LiAc, with the volume adjusted to supply 10 μl of cell suspension per transformation. For each transformation, 10 μl of the prepared suspension was transferred to a PCR tube and centrifuge. After removing the supernatant, the cell pellet was resuspended in 10 μl of transforming DNA mixture. Then, 72 μl of a LiAc/SS carrier DNA/PEG solution—comprising 53 μl of sterile 50% (w/v) PEG-3350 (Sigma), 8 μl of sterile 1 M LiAc and 11 μl of single-stranded carrier DNA (salmon sperm DNA, 2mg/ml)—was added and mixed thoroughly. The transformation mix was incubated at 42°C for 30 minutes and 30°C for 30 minutes. Then the cells were pelleted by centrifugation, the supernatant was discarded, and the pellet was resuspended in 100 μl of sterilized water. The resulting suspension was plated onto YNB+N-U plates for auxotrophic selection and incubated at 30°C for 2 days until colonies formed.

### Growth and condition screens of co-culture starters

Synthetic starter media were prepared by mixing synthetic flour powder (90% potato starch [Sigma] and 10% wheat gluten [Sigma]) with various growth media. Bob’s Red Mill whole wheat flour was used to demonstrate that starter media could be constituted from commercial flour. The flour powder was incubated at 95°C for 1 hour to sterilize. *L. plantarum* WCFS1 or NC8 and *S. cerevisiae* BY4741 MATa were inoculated into MRS and YPD broth for overnight culture. Overnight cultures of *L. plantarum* and *S. cerevisiae* were inoculated 1:100 into fresh MRS and YPD medium, grown to early log phase (OD_600_ between 0.3 and 0.5), then diluted to a final concentration of 2×10^5^ colony-forming units (CFU) ml^−1^ in PBS. 2 ml liquid medium and 50 μl of diluted *L. plantarum* and *S. cerevisiae* were added to 2 g of synthetic flour powder in 14 ml round-bottom test tubes and homogenized. Co-cultures were incubated at 30°C under static conditions. To quantify the growth of *L. plantarum* and *S. cerevisiae*, 10 μl of homogenized co-culture products were harvested at 48 hours, and 10-fold serial dilutions were plated on both MRS and YPD agar plates. CFU per unit volume were counted.

For daily transfer, the synthetic starter cultures were homogenized by stirring, and 100 μl of the 24-hour co-culture products from the previous day was used to inoculate fresh starter media. Three rounds of serial transfers were conducted for each sample.

To inoculate *L. plantarum* and *S. cerevisiae* at ratios of 1:1, 100:1, and 10,000:1, 50 μl of *S. cerevisiae* (2×10^5^ CFU ml^−1^ in PBS) and 50 μl of *L. plantarum* at cell concentrations of 2×10^5^, 2×10^7^, and 2×10^9^ CFU ml^−1^ in PBS were added to 2 ml MRS broth + 2 g of synthetic flour.

### Scanning electron microscopy (SEM)

Synthetic starters grown for 24 hours were freeze-dried with a lyophilizer. The dried samples were coated with a gold sputter, and images were taken with a JEOL-6390 scanning electron microscope.

### Transcriptomic analysis of synthetic starter

For transcriptomic comparison of co-culture and mono-culture systems, three sets of synthetic starters were constructed: co-culture, *L. plantarum* mono-culture, and *S. cerevisiae* mono-culture. To extract microbes from starters, ice-cold RNA stop solution (5% water-saturated phenol in ethanol) was added to the starter cultured for 24 hours, followed by ice-cold PBS and mixing. *L. plantarum* was extracted by low-speed centrifugation (200 rcf) at 4°C for 10 minutes; the supernatant, containing *L. plantarum* cells, was collected and washed with PBS. The sediment of centrifugation was resuspended with ice-cold PBS, allowing for 15 minutes of static precipitation, and the supernatant, containing *S. cerevisiae* cells, was collected. *L. plantarum* and *S. cerevisiae* pellets were lysed enzymatically, with lysozyme (Thermo) and zymolyase (Zymo Research), respectively. Total RNA of each sample was extracted using a QIAGEN RNeasy Mini kit following the manufacturer’s protocol.

For the analysis of RNA-seq data, Trim Galore was first used to process the raw sequencing reads by quality control and removing adaptors. Then, HISAT2 was used to align the reads to the reference genome of *L. plantarum* WCFS1 and *S. cerevisiae* BY4741 (*64*), followed by generating an expression matrix with featureCounts (*65*). DEseq2 was used for differential analysis (*66*), and gene set enrichment results were analyzed with R package ClusterProfiler (*67*).

### Metabolomics of amino acids in synthetic starter

Mono-cultures of *L. plantarum* and *S. cerevisiae*, co-cultures consisting of both species, and synthetic starter media alone were homogenized in ice-cold water and freeze-dried prior to metabolomic analysis. For sample preparation, 80 mg of each sample was combined with 400 μl of a cold methanol/acetonitrile solution (1:1, v/v) containing internal isotope standards. Lysates were homogenized using an MP homogenizer (20 seconds, three times), vortexed, sonicated for 5 minutes at 4°C, then incubated at −20°C for 1 hour. The mixture was centrifuged for 20 minutes (14000 rcf, 4°C). The supernatant was dried in a vacuum centrifuge, and the samples were re-dissolved in 150 μl acetonitrile/water (1:1, v/v), vortexed, and centrifuged (14000 rcf, 4°C, 15 min). Supernatants were collected for LC-MS/MS analysis.

The samples were separated using an Agilent 1290 Infinity LC ultra-high performance liquid chromatography system (UHPLC) with a hydrophilic interaction liquid chromatography (HILIC) column at a column temperature of 35°C, a flow rate of 0.3 ml min^-1^, an injection volume of 2 μl, and a mobile phase composition of A: 90% water + 2 mM ammonium formate + 10% acetonitrile, and B: acetonitrile + 0.4% formic acid. The gradient elution program was as follows: 0—1.0 min, 85% B; 1.0—3.0 min, B linearly changed from 85% to 80%; 3.0—4.0 min, 80% B; 4.0—6.0 min, B linearly changed from 80% to 70%; 6.0—10.0 min, B linearly changed from 70% to 50%; 10—15.5 min, B maintained at 50%; 15.5—15.6 min, B linearly changed from 50% to 85%; 15.6—23 min, B maintained at 85%. To prevent fluctuations in instrument detection signals and avoid their impact, the samples were kept in an automatic sampler at 4°C and analyzed in a random order. Quality control (QC) samples were inserted into the sample queue to monitor and evaluate the stability of the system and ensure the reliability of the experimental data. Mass spectrometry analysis was performed using an AB 6500+ QTRAP mass spectrometer (AB SCIEX). The ESI source conditions were as follows: Source temperature: 580°C, Ion Source Gas1 (GS1): 45 psi, Ion Source Gas2 (GS2): 60 psi, Curtain Gas (CUR): 35 psi, IonSpray Voltage (IS): +4500 V or −4500 V in positive or negative modes, respectively, using MRM mode for monitoring.

### Amylase assay

Overnight cultures of engineered *L. plantarum* or *S. cerevisiae* candidate strains were inoculated at a ratio of 1:100 into fresh medium and cultured for 24 hours. The supernatant was harvested and processed for amylase activity measurement. The amylase secretion capacity of *L. plantarum* strains was determined with the Phadebas Amylase test kit, following the manufacturer’s protocol. To determine the texture-changing capability in the synthetic starter system, the starters were prepared as previously described and inoculated with different combinations of *L. plantarum* and *S. cerevisiae* strains. Heights were measured after 24 hours of culturing.

### Rheology test

Starters were grown from wild-type yeast with wild-type or yLF201 amylase-secreting *L. plantarum*. Rheological measurements were conducted using around 50 μl of homogenized starters on a TA Instruments ARES-G2 strain-controlled rheometer. The rheometer was equipped with a customized steel parallel plate geometry, with the upper fixture measuring 8 mm in diameter and the bottom fixture measuring 25 mm in diameter. All measurements were performed at room temperature. During the frequency sweep tests, the frequency varied from 100 to 0.01 rad/s while a constant strain of 0.1% was maintained.

### Characterizing *L. plantarum* sensing strains

To characterize the *L. plantarum* sensor strains, overnight cultures were inoculated at a ratio of 1:100 into fresh medium and cultured for 3 hours to early log phase. Then, the corresponding ligand (DAPG or IP-673) with the desired concentration was supplemented. After 18 hours of induction, the cells were harvested for fluorescence or luciferase measurements. For fluorescence measurement, the cells were washed and resuspended with the same volume of PBS, then transferred to a black 96-well plate for fluorescence measurement with a microplate reader (Agilent BioTek Synergy H1). (Washing and resuspension with PBS removed the MRS supernatant, which would have increased background during the measurement of mCherry fluorescence.) For the NanoLuc luciferase assay, the Nano-Glo luciferase assay system (Promega, N1120) was used. 100 μl of culture was combined with the same volume of assay mix, and the mixture was incubated for 5 minutes, followed by microplate readout. The dose-response curves were fitted with the R drc package using a four-parameter logistic model. To characterize the OR-gate, 100 ng ml^-1^ of IP-673 or 10^4^ nM of DAPG was used for activation in designated situations. To characterize the OR-gate sensor in the synthetic starter, the Design 2 OR-gate *L. plantarum* and wild-type *S. cerevisiae* were inoculated as stated previously and cultured in black 96-well plates with an adjusted media volume of 1.5 g synthetic flour + 1.5 ml MRS broth. IP-673 or DAPG was added to the starter upon inoculation. The starters were harvested for luciferase measurement after 24 hours of culture. For the luciferase assay, 750 μl of Nano-Glo assay mix was added to the synthetic starter in 12-well black plates. The mix was well-homogenized and incubated for 5 minutes in the dark, followed by luminescence visualization on a Biorad Chemidoc MP system.

### Characterizing communication systems

To characterize the IP-673-based communication system, overnight cultures of *L. plantarum* receiver strains (swF115 and swF221, for IP-673 inducible fluorescence and NanoLuc luciferase output, respectively) and *S. cerevisiae* sender strains (constitutively secreting IP-673) were inoculated at a ratio of 1:100 into fresh MRS and YPD medium and allowed to grow to an OD_600_ between 0.3 and 0.5, and then diluted to a final concentration of 2×10^5^ CFU ml^−1^ in fresh MRS medium. In each well of a 96-well plate, a total of 10000 *L. plantarum* and *S. cerevisiae* cells in PBS were inoculated into 190 μl fresh MRS, with different ratios of sender and receiver strains versus control strains. After incubating for 24 hours statically at 30°C, the cultures were harvested and washed with PBS; fluorescence or luciferase was then measured. To characterize the ability of consortia to sense various stimuli, the *L. plantarum* receiver strains and sensor-sender *S. cerevisiae* (secreting IP-673 when induced by corresponding signals) were prepared in the same way, with corresponding signals administered.

To characterize the communication system in the synthetic starter, *S. cerevisiae* constitutive sender or sensor-sender strains and *L. plantarum* swF221 receiver strains were inoculated as stated previously and grown in black 96-well plates with an adjusted media volume. For sensor-sender cases, inducing molecules with designated concentrations for activating the yeast sensor were added, upon inoculation, to the MRS broth that was used for making synthetic starter. Starters were harvested for luciferase measurement after 24 hours of culture. For the luciferase assay of the synthetic starter, 750 μl of Nano-Glo assay mix was added to the synthetic starter in 12-well black plates; the starter was grown from 1.5 g of synthetic flour and 1.5 ml MRS medium for BED- and MFα-sensing consortia, and 250 μl assay mix was added to the synthetic starter grown from 0.5 g flour and 0.5 ml MRS for characterizing blue-light-sensing consortia. The mix in each case was well-homogenized and incubated for 5 minutes in the dark, followed by luminescence visualization on a BioRad Chemidoc MP system.

### Animals

Male C57BL/6J mice aged 6-8 weeks were procured from the Chinese University of Hong Kong (Hong Kong SAR, China) and housed in standard enclosures within a controlled environment, at 23 ± 2°C and a 12-hour light/dark cycle, with *ad libitum* access to food and water. All procedures for handling the mice and conducting the experiments were carried out in accordance with the guidelines set forth by the Committee on the Use of Human and Animal Subjects in Teaching and Research (HASC) of Hong Kong Baptist University (HASC registration# REC/19-20/0301). The study followed the regulations stipulated in the Animal Ordinance of the Department of Health, Hong Kong SAR, China. For the DSS-induced colitis mouse model, following a period of acclimatization feeding lasting 1 week, colitis was induced by administering a 2% dextran sulfate sodium (DSS) solution (MP Biomedicines, USA) in the drinking water for 5 days, followed by a 7-day recovery.

### *In vivo* delivery of probiotic consortia of synthetic starter

*L. plantarum* constitutively expressing mCherry fluorescence and *S. cerevisiae* expressing mTurquoise2 were inoculated into the liquid culture or synthetic starter. 0.2 ml of synthetic starter and 0.2 ml PBS suspension of *S. cerevisiae* and *L. plantarum* with approximately the same numbers as in synthetic starter were used for gavage. Feces were collected at designated time points, suspended with PBS (1 ml PBS for 100 mg feces) and homogenized with Qiagen Tissue Lyser, followed by serial dilution and plating on MRS plates with 10 µg ml^−1^ erythromycin and on YNB+N-U plates with chloramphenicol 34 µg ml^−1^ and kanamycin 50 µg ml^−1^ for CFU counting. Colonies were double-checked for fluorescence. Final cell counts were calculated by normalizing according to the original cell counts in the PBS mix and according to the SINERGY cell counts used for gavage. CFU values below the detection limit of plating were plotted and included in statistical analysis as zero for accessible visualization and calculation.

### Visualizing SINERGY consortia *in vivo*

Mice fasting for 12 hours were administered with 0.2 ml of synthetic starter with *L. plantarum* constitutively expressing mCherry fluorescence and *S. cerevisiae* expressing super-fold GFP (sfGFP). Mice were sacrificed 3 hours after gavage, and proximal colons were harvested. The harvested tissues were promptly placed in a 4% paraformaldehyde solution in PBS for fixation for 24 hours. Subsequently, the tissue was dried in a series of sucrose solutions (15% for 24 hours and 30% for 24 hours), then embedded in O.C.T. compound and sectioned to a 10 μm thickness on a Thermo Scientific CryoStar NX70. The slices were stained with 4’,6-diamidino-2-phenylindole (DAPI), followed by a PBS wash and mounting in ProLong™ Gold Antifade Mountant (Thermo Fisher Scientific). Imaging was performed with a Leica DMi8 Confocal Laser Scanning Microscope.

### Transposon mutagenesis of *L. plantarum*

Wild-type *L. plantarum* NC8 was first transformed with a plasmid harboring constitutively expressed *orf1/2* genes (putative transposases of IS1223) under the strong promoter P_tlpA_ (*68*), with a thermo-sensitive origin of replication from pVE6007 (*69*). A suicide transposon donor that lacks a *Lactobacillus* origin of replication and carries the engineered IS1223 transposon was then electroporated into the previous strains. The engineered transposon includes a chloramphenicol resistance gene and an mCherry reporter expressed under the P_11_ promoter(*70*), flanked by inverted repeat sequences from IS1223. After electroporation, chloramphenicol-resistant colonies were collected. A previous study demonstrated that methylation patterns of plasmid DNA dramatically influence the electroporation efficiency of *L. plantarum*(*71*). To promote electroporation efficiency, non-methylated suicide plasmids harvested from NEB C2925 competent *E. coli* (dam–/dcm–) were used for electroporation. 5 μl of suicide plasmid (in total 5 μg) was added to 95 μl of *L. plantarum* NC8 competent cells. Multiple rounds of electroporation were conducted to generate sufficient numbers of mutants, followed by selection on MRS agar with 10 μg ml^-1^ chloramphenicol for 48 hours. Around 4000 colonies were collected as a pooled Tn-mutant library. Colony PCR was used to verify integration using the primers. Arbitrary PCR (abPCR) was used for the small-scale check of insertion loci using the protocols described by Saavedra et al. (*72*) The primers used for colony PCR and abPCR are available in **Table S1**.

### Targeted enrichment sequencing and mapping of insertion sites

Genomic DNA of the Tn library was extracted with lysozyme lysis and QIAGEN DNeasy Blood & Tissue Kits. NGS libraries were first prepared using the Homgen Universal DNA Library Prep Kit. Transposon boundary sequences were used to design 120-bp probes for the targeted enrichment of sequences that contain the boundary of genomic insertion (**Table S1**). Hybridization capture was conducted using the Homgen ProbeCap system, followed by PE300 sequencing. For processing sequencing data, trimmed sequencing reads were mapped to both the genomic sequence of *L. plantarum* NC8 and the probes using BWA and Samtools. Only the reads that mapped to both the genome and the probe sequence were kept for precise determination of the insertion sites. The insertion sites were then mapped to genomic features and visualized by Proksee (*73*).

### Screening of Tn mutants and INSeq analysis

For *in vitro* screening, overnight cultures of Tn mutants were diluted into fresh MRS with a 1:50 ratio and grown to an OD_600_ = 0.4. The bacteria were then washed with PBS and treated with designated sodium cholate (0.2%, 0.4% or 0.8%) or pH conditions (pH = 2, 3 or 3.5) achieved by appropriate concentrations of HCl, followed by plating on MRS plate with 10 µg ml^−1^ chloramphenicol. Surviving cells were scraped, cultured, and treated again for a total of 3 rounds under corresponding conditions, with three orthogonal groups proceeding separately. After final treatment, the genomic DNA of the libraries was extracted and sequenced to identify insertion sites. For *in vivo* screening, 0.2 ml of synthetic sourdough starter with Tn-mutant libraries was administered into mice by gavage. Mutants that resided for 24 hours were collected by plating feces, and the screening was repeated for 3 rounds. Three groups of selections were conducted separately. Insertion sites were sequenced and mapped as previously described. DEseq2 was used for differential analysis to identify mutants that increase or decrease fitness in each condition. Insertions with an adjusted p-value < 0.05 and a log_2_[foldchange] > 1 were marked as increased-fitness insertions; those with an adjusted p-value < 0.05 and a log_2_[foldchange] < −1 were marked as decreased-fitness insertions. Other insertions were marked as non-significant insertions (NS).

### Construction and characterization of GPCR-based *S. cerevisiae* sensor

All yeast GPCR-based sensor strains were constructed based on the yWS677 strain and yeast MoClo toolkits (*54*, *74*). For the eATP sensor, the P2Y2 receptor sequences of the five organisms included were obtained from Uniprot and codon-optimized for *S. cerevisiae* expression. Overnight cultures of yeast sensor strains were diluted 100-fold into fresh YNB+N-U (pH = 7, buffered by 50 mM potassium phosphate buffer) with designated concentrations of ATP (pH = 7), and cultured for 18 hours. The cells were washed and resuspended with the same volume of PBS before GFP fluorescence measurement. Dose-response curves were generated with an R drc package (*75*). The parameters of the sensor strains (tightness, dynamic range, sensitivity, operational range, and maximum output) were defined as described by Shaw et al. (*54*). For the niacin sensor, human HCA2 sequences were obtained from Uniprot and codon-optimized. Measurements were the same as used for the eATP ones, with designated concentrations of niacin added.

### Characterization of GPCR-based SINERGY consortia

For *in vitro* characterization of the eATP-sensing consortia, saturated cultures consisting of swF115 (IP-673 inducible mCherry expression) *L. plantarum* strain with Design 7 or Design 10 eATP sensor *S. cerevisiae* strain in MRS broth (pH = 7) were induced with designated concentrations of ATP. The niacin-sensing consortia consisted of the swF115 (IP-673 inducible mCherry expression) *L. plantarum* strain and the swY035 niacin sensor *S. cerevisiae* strain (Gpa chimera expressed by promoter P_GPK1_) in MRS broth (pH = 7), induced with designated concentrations of niacin. Fluorescence levels 6 hours after the induction were measured by a microplate reader.

For *in vivo* characterization of the eATP-sensing consortia, 0.2 ml of SINERGY with the Design 10 eATP sensor *S. cerevisiae* and swF221 (IP-673 inducible NanoLuc secretion) *L. plantarum* were administered into mice with colitis induced by 2% DSS (n = 9) on day 7. Day 7 was the expected peak of the disease, when increased ATP levels were anticipated in the intestinal tract, with healthy mice as the control (n = 6). The feces were collected 6 hours after gavage, homogenized in buffered MRS medium (pH = 7), and incubated for 12 hours before measuring NanoLuc luminescence levels in the culture.

### Evaluation of disease activity index (DAI)

To assess colitis severity, mice were evaluated using the Disease Activity Index (DAI). The DAI score was determined by summing the scores for weight loss (0, <1%; 1, 1%-5%; 2, 5%-10%; 3, 10%-15%; 4, >15%), the presence of diarrhea (0, normal stools; 1, soft and shaped stools; 2, loose stools; 3, liquid stool; 4, diarrhea), and fecal occult blood detection (0, none blood occurrence; 1, weak positive bleeding; 2, positive bleeding; 3, strong positive bleeding; 4, gross bleeding) (*76*).

### Quantification of therapeutic payloads secreted by engineered *L. plantarum*

Therapeutic payloads (murine IL-10, human Elafin, murine SLPI) were codon-optimized for *L. plantarum* expression. *L. plantarum* strains with payloads and various signal peptides were induced at OD=0.3 with 100 ng ml^−1^ IP-673, and supernatant was harvested after 5 hours of induction in buffered MRS (pH=7). ELISA was conducted to quantify the amount of payload in the culture supernatant, using a Mouse IL-10 Uncoated ELISA Kit (Invitrogen), a Human Trappin-2 / Elafin ELISA Kit (Merck), and a Mouse SLPI ELISA Kit (UpingBio), respectively, following the manufacturer’s protocol.

### DSS-treated mice administered with SINERGY

Mice in the DSS-WT and DSS-TH groups were administered daily with 0.2 ml of homogenized wild-type or therapeutic SINERGY via gavage, starting on day 0 of the experiment. Therapeutic SINERGY consisted of approximately 5 × 10^9^ CFU *L. plantarum* (equal numbers of strains secreting sp3050-mIL-10 and sp0297-hElafin were inoculated) and 5 × 10^7^ CFU of *S. cerevisiae*. In the DSS-PBS group, 0.1 ml PBS was administered daily, due to the ∼50% water content of SINERGY.

### Histopathological analysis

Distal colon tissues were harvested and fixed with 4% paraformaldehyde solution. Subsequently, the tissue was dried using a series of sucrose solutions (15% for 24 hours and 30% for 24 hours), followed by embedding in O.C.T. compound and sectioning to a 10 μm thickness on Thermo Scientific CryoStar NX70. Sections were stained with hematoxylin and eosin (H&E) (G1120, Solarbio, Beijing, China) and observed with an optical microscope (Leica, Wetzlar, Germany). Histological damage to the colonic tissue was assessed based on scoring for inflammation cell infiltration (0, non-infiltration; 1, mucosal infiltration; 2, mucosal and submucosal infiltrate of inflammatory cells; 3, mucosal and submucosal infiltration with crypt abscesses; 4, transmural inflammatory cells) and epithelial changes (0, normal architecture; 1, basal 1/3 crypt damage or minimal goblet cell loss; 2, basal 2/3 crypt damaged or goblet cell loss; 3, ulceration with only surface epithelium intact, 4, epithelium loss).

### Analysis of cytokines in colon tissues

Following sacrifice of the mice, the colon tissues were harvested and homogenized in RIPA lysis buffer (every 10 mg in 100 μl) and centrifuged for supernatant collection. The homogenates were processed with LEGENDplex MU Inflam Panel (Biolegend, USA) and measured with BD FACSCelesta following the manufacturer’s protocol. Values below the detection limit were plotted and included in statistical analysis as the detection limit value for accessible visualization and calculation.

### Microbiome analysis of mouse feces

On Day 11 of the experiment, the feces from each mouse were collected. Total genomic DNA was extracted from the samples using the Mag-bind Soil DNA Kit (Omega), and its purity and concentration were assessed. The V3-V4 variable region was selected for sequencing. Specific primers with barcodes and high-fidelity DNA polymerase were used to amplify this region via PCR. The PCR products were visualized using 2% agarose gel electrophoresis, and the desired fragments were recovered using the Quant-iT PicoGreen dsDNA Assay Kit. The library was prepared using the TruSeq Nano DNA LT Library Prep Kit from Illumina. The constructed library was assessed for quality using the Agilent Bioanalyzer 2100 and Promega QuantiFluor. The data was analyzed with QIIME2 (*77*).

### Metabolomics of mouse intestinal contents

The mice’s cecal contents were harvested upon sacrifice and stored at −80°C. For sample preparation, the samples were thawed at 4°C, and 80 mg of each sample was combined with 400 μl of methanol/acetonitrile/water (2:2:1, v/v/v) and homogenized. Homogenized samples were sonicated for 30 minutes at low temperature, then incubated at −20°C for 10 mins. The mixture was centrifuged for 20 minutes (14000 rcf, 4°C). The supernatant was dried in a vacuum centrifuge, the dried samples re-dissolved in 100 μl acetonitrile/water (1:1, v/v), vortexed, and then centrifuged (14000 rcf, 4°C, 15 min). Supernatants were collected for LC-MS/MS analysis.

The LC-MS/MS Analysis was carried out using a Vanquish UHPLC system coupled to an Orbitrap Exploris 480. For hydrophilic interaction liquid chromatography (HILIC) separation, samples were run on a 2.1 mm × 100 mm ACQUIY UPLC BEH Amide 1.7 μm column (Waters, Ireland). The mobile phase in both electrospray ionization (ESI) positive and negative modes consisted of A = 25 mM ammonium acetate and 25 mM ammonium hydroxide in water, and B = acetonitrile. The gradient started at 95% B for 0.5 minutes, then linearly decreased to 65% over 6.5 minutes, further reduced to 40% in 1 minute and held for 1 minute, and finally increased back to 95% in 0.1 minute and maintained for 2.9 minutes. The ESI source conditions were as follows: Gas1 at 50 psi, Gas2 at 2 psi, source temperature at 350°C, and IonSpray Voltage Floating (ISVF) at +3500 V/-2800V. In the MS-only acquisition mode, the instrument was set to scan the m/z range of 70-1200 Da, with a resolution of 60000 and an accumulation time of 100 ms. For auto MS/MS acquisition, the instrument scanned the m/z range of 70-1200 Da, with a resolution of 60000, an accumulation time of 100 ms, and exclusion within 4 seconds.

### Quantification and statistical analysis

Replicates and details of statistical analysis were used in all experiments, as noted in figure captions or methods. Unpaired two-sided t-test or Mann-Whitney test was applied for comparisons between two groups. Two-way analysis of variance (ANOVA) was applied for multiple group comparisons with time-course data. Graphpad Prism 9 was used for data statistical analysis. The flow cytometry data from LEGENDplex MU Inflam Panel were measured with BD FACSCelesta, with at least 4000 events per sample for the 13-plex assay, as guided by the manufacturer’s protocol. The LEGENDplex Data Analysis Software Suite was used for initial quality control checks. The automated gating of bead population was reviewed and manually adjusted for any misclassification. All graphs and data visualizations were generated using R unless specified otherwise.

## Acknowledgments

We thank Karen Pepper for editing the manuscript and Elizabeth Landis for advice and discussions. We thank Tom Ellis for generously providing the yWS677, yWS890 strains and GPH093 plasmid. We thank Fei Sun and Qikun Yi for assisting with rheological testing.

## Author contributions

Conceptualization: T-C.T. and Y.L.;

Methodology: S.W., L.C., W.M.S., W.J.X., H.Y.T., C.L., X.J., T-C.T., and Y.L.;

Investigation: S.W., Y.X., Y.Z., M.E.G., S.C.Y.C., L.C., T-C.T., and Y.L.;

Visualization: S.W. and Y.L.;

Funding acquisition: ZX.B. and Y.L.;

Supervision: H.Y.T., ZX.B., P.D., G.M.C., T.K.L., T-C.T., and Y.L.;

Writing – original draft: S.W. and Y.L.;

Writing – review & editing: S.W., T-C.T., and Y.L with input from all authors.

## Data and materials availability

Raw sequencing data for transcriptomic analysis of the synthetic starter have been deposited at the NCBI SRA database under accession no. PRJNA1308853. Targeted enrichment sequencing data of *L. plantarum* Tn-mutants have been deposited at the NCBI SRA database under accession no. PRJNA1309084. Raw sequencing data for 16S microbiome analysis of mouse feces have been deposited at the NCBI SRA database under accession no. PRJNA1309613. This study does not report original code. The following plasmids will be made available through Addgene: ylF167 (DAPG-inducible design 2 plasmid), swF179 (constitutive expression of IS1223 transposases), and swF238 (suicide donor of engineered IS1223 transposon).

