## Supplementary Figures for "Sourdough starter-inspired living materials grown from probiotic consortia with engineered division-of-labor"

Shuchen Wang *et al.*

**The PDF file includes:**

Figs. S1 to S10

**Other Supplementary Materials for this manuscript includes the following:**

Table S1.

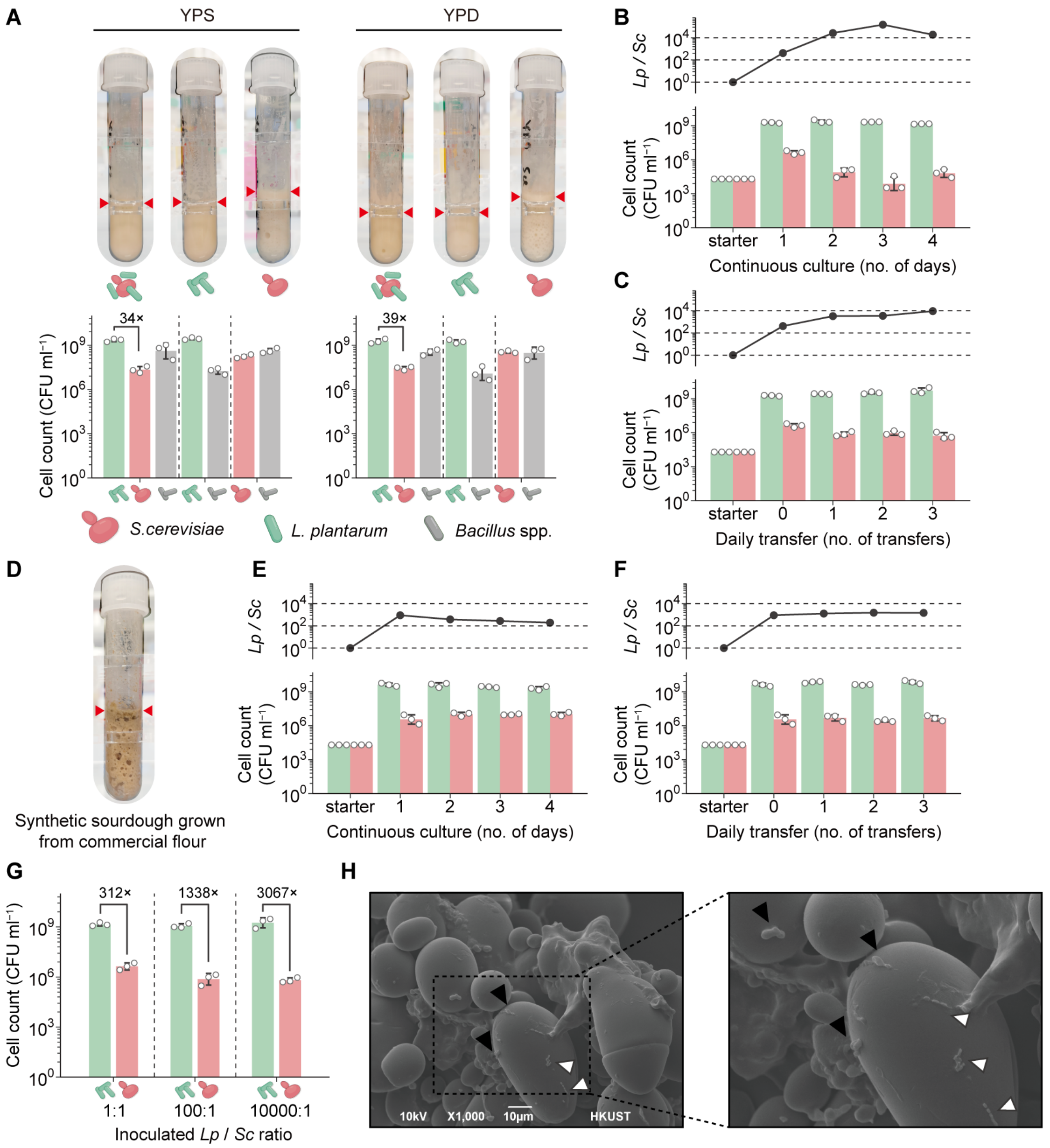


**Fig. S1: Inoculation conditions to determine the texture and composition of synthetic starter.**

(**A**) Cell counts and images of synthetic starter grown from YPD and YPS (mean ± sd, n = 3 starters). The red triangular arrows indicate the fermentation height of the synthetic starter. Grey bars indicate uninoculated *Bacillus* spp. in the starter, likely originating from the gluten product.

(**B** and **C**) Cell counts and ratio of *L. plantarum* NC8 to *S. cerevisiae* (*Lp* / *Sc*) over days of continuous culture (**B**) and daily transfer (**C**) (cell count represented by mean ± sd, ratio represented by mean value, n = 3 starters).

(**D**) Image of synthetic sourdough starter grown from commercial flour.

(**E** and **F**) Cell counts and *Lp* / *Sc* ratio over days of continuous culture (**E**) and daily transfer (**F**) with commercial flour (cell counts represented by mean ± sd, ratio represented by mean value, n = 3 starters).

(**G**) Cell counts of the synthetic starter grown from different inoculation ratios (cell counts

represented by mean ± sd, n = 3 starters).
(**H**) Representative SEM imaging of the synthetic starter. Black arrow: *S. cerevisiae* cells; white arrow: *L. plantarum*.

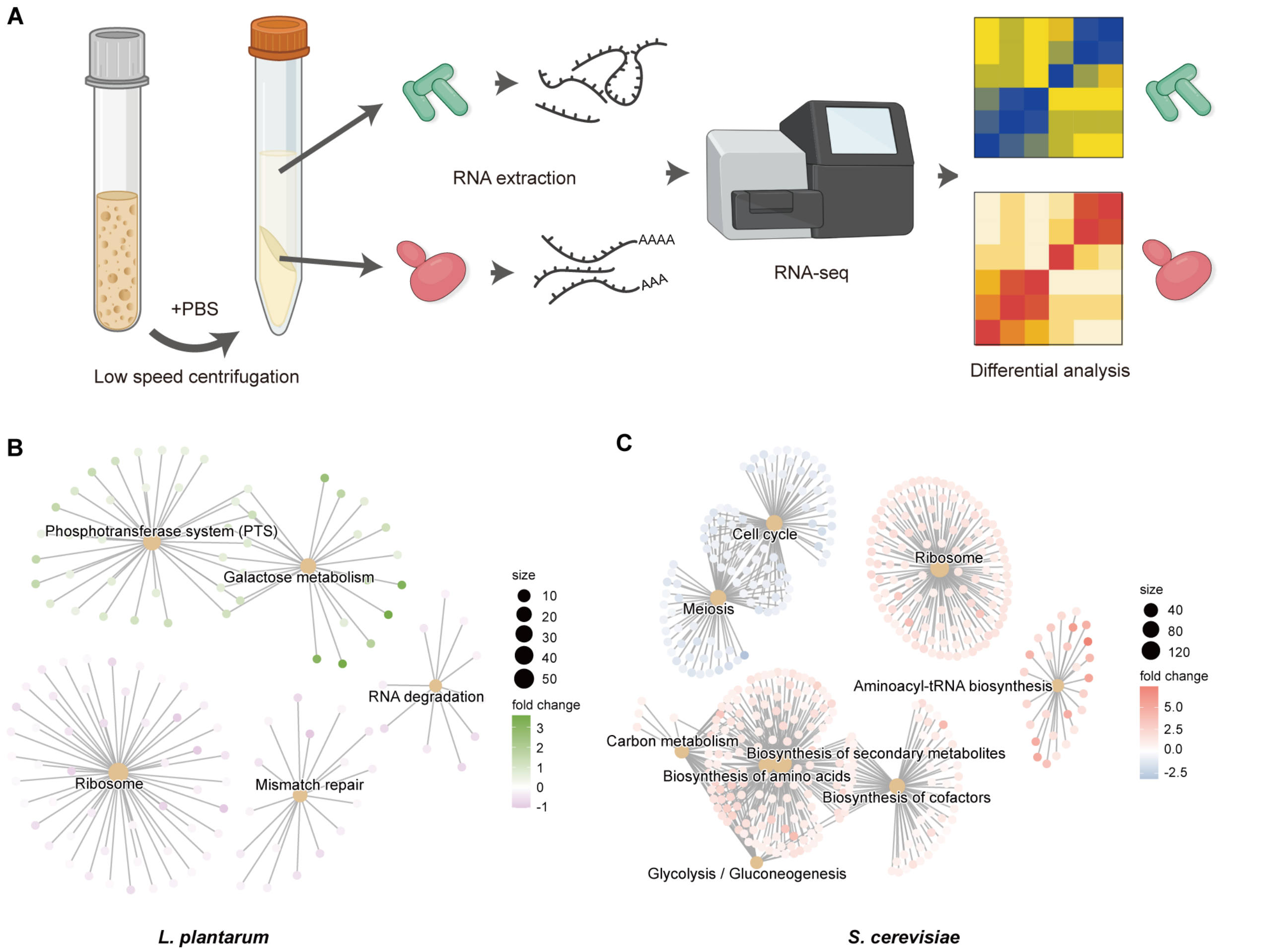


**Fig. S2: Transcriptomic analysis to reveal molecular interactions between *L. plantarum* and *S. cerevisiae* in synthetic starter.**

(**A**) Schematic of transcriptomic analysis workflow.

(**B** and **C**) Gene set enrichment assay of RNA-seq results for *L. plantarum* (**B**) and *S. cerevisiae* (**C**) under co-culture conditions compared with mono-culture conditions.

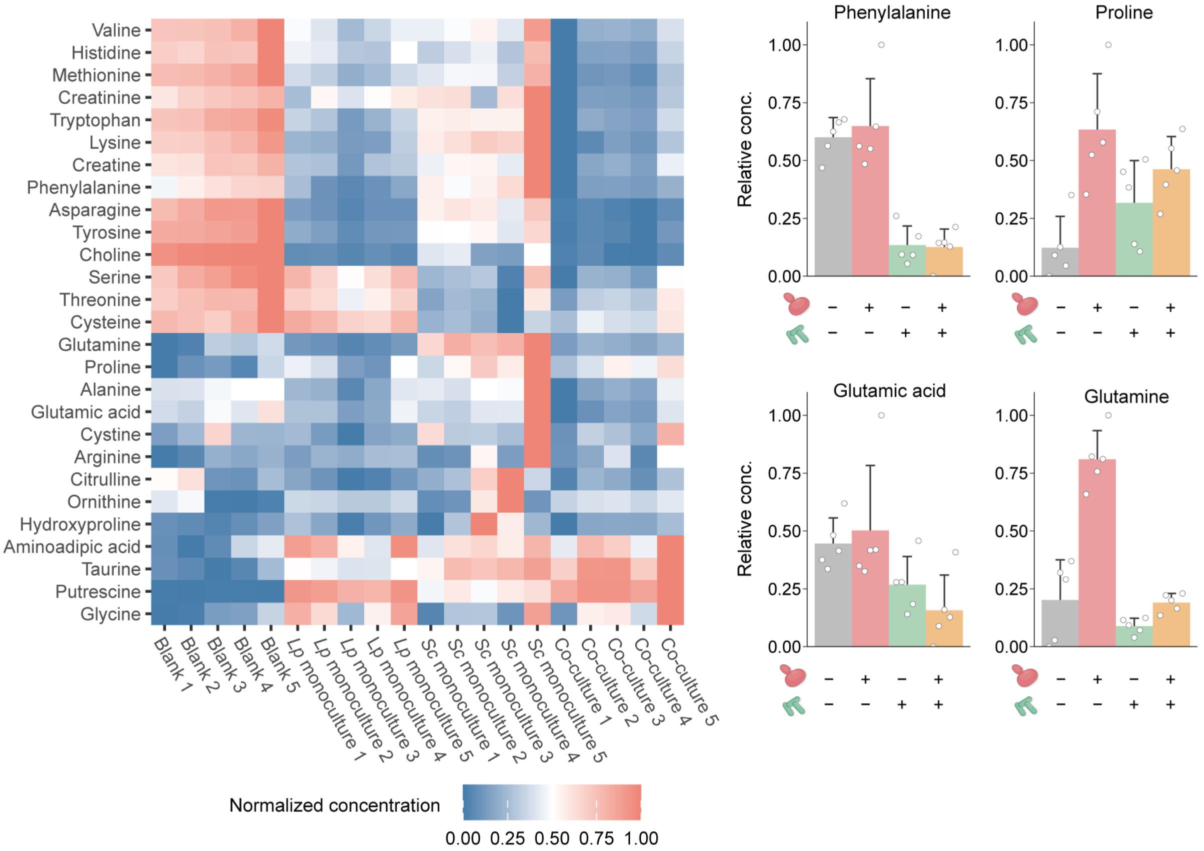


**Fig. S3: Metabolomic analysis of SINERGY.**

Targeted metabolomics of amino acids in synthetic starter from blank medium, *L. plantarum* mono-culture, *S. cerevisiae* mono-culture, and co-culture (mean ± sd, n = 5 starters).

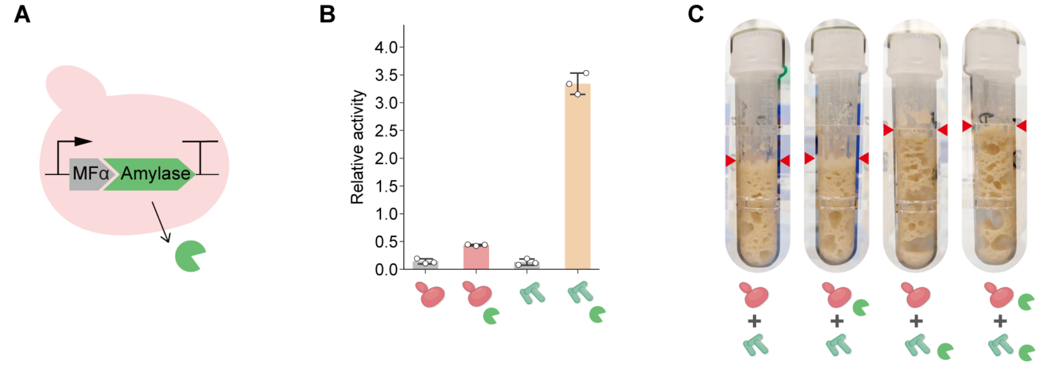


**Fig. S4: Characterizing amylase-secreting *S. cerevisiae*.**
(**A**) Schematic of the engineered amylase-secreting *S. cerevisiae* strain.
(**B**) Characterization of amylase-secreting capacity of the *S. cerevisiae* strain and comparison with *S. cerevisiae* wild-type strain, *L. plantarum* wild-type, and the ylF201 amylase-secreting *L. plantarum* strains (mean ± sd, n = 3 independent cultures).
(**C**) Representative images from the combinatorial co-culturing of *L. plantarum* and *S. cerevisiae* wild-type or amylase-secreting strain.

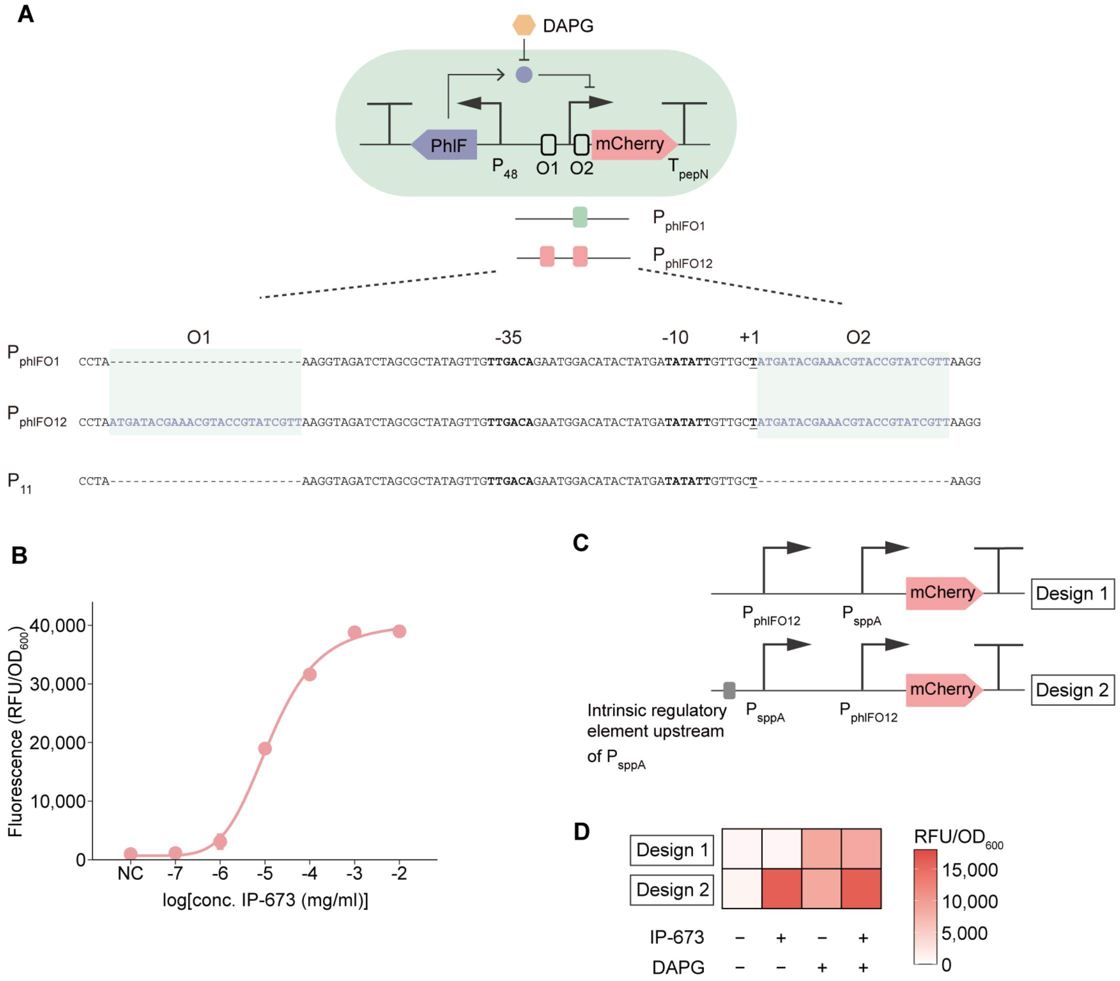


**Fig. S5: Design and characterization of DAPG/IP-673 OR-gate in *L. plantarum*.**
(**A**) Design of PhlF-binding promoters. PhlF operator sites were inserted in various regions (O1 or O2) of the $P_{11}$ promoter.
(**B**) Dose-response properties of IP-673 peptide-inducible system in *L. plantarum* (mean ± sd, n = 3 independent cultures).
(**C**) Design of DAPG/IP-673 OR-gate *L. plantarum* strains.
(**D**) Characterization of the OR-gate in MRS broth with fluorescence output (mean, n = 3 independent cultures).

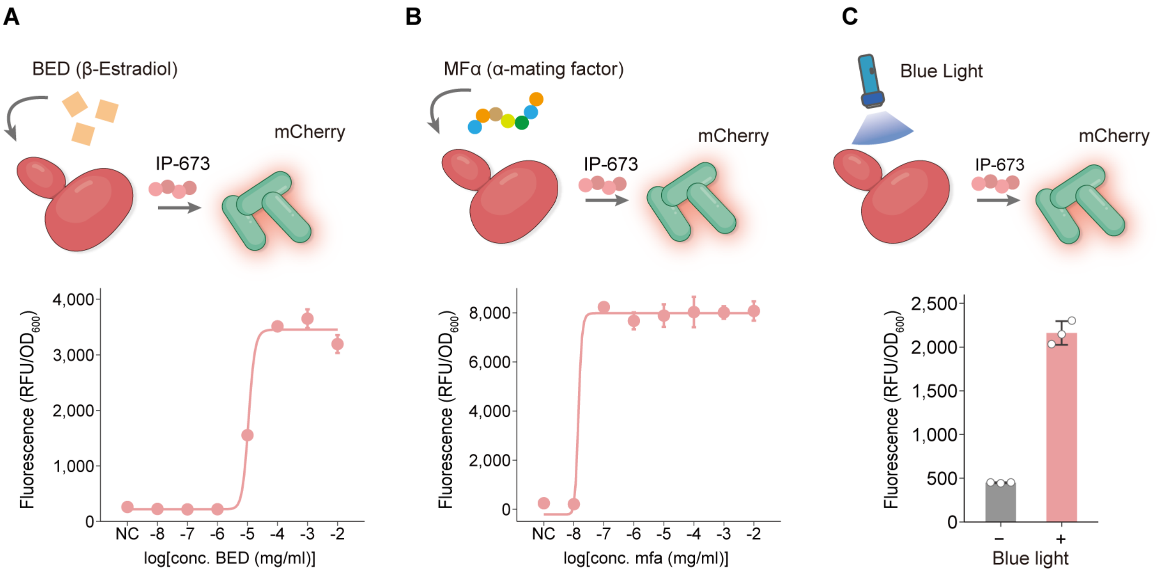


**Fig. S6: Characterization of the communication system with mCherry fluorescence as reporter.**
(**A**-**C**) The responses of engineered consortia to various signals – (**A**) BED, (**B**) MFα (**C**) Blue light (mean ± sd, n = 3 independent cultures). NC, negative control.

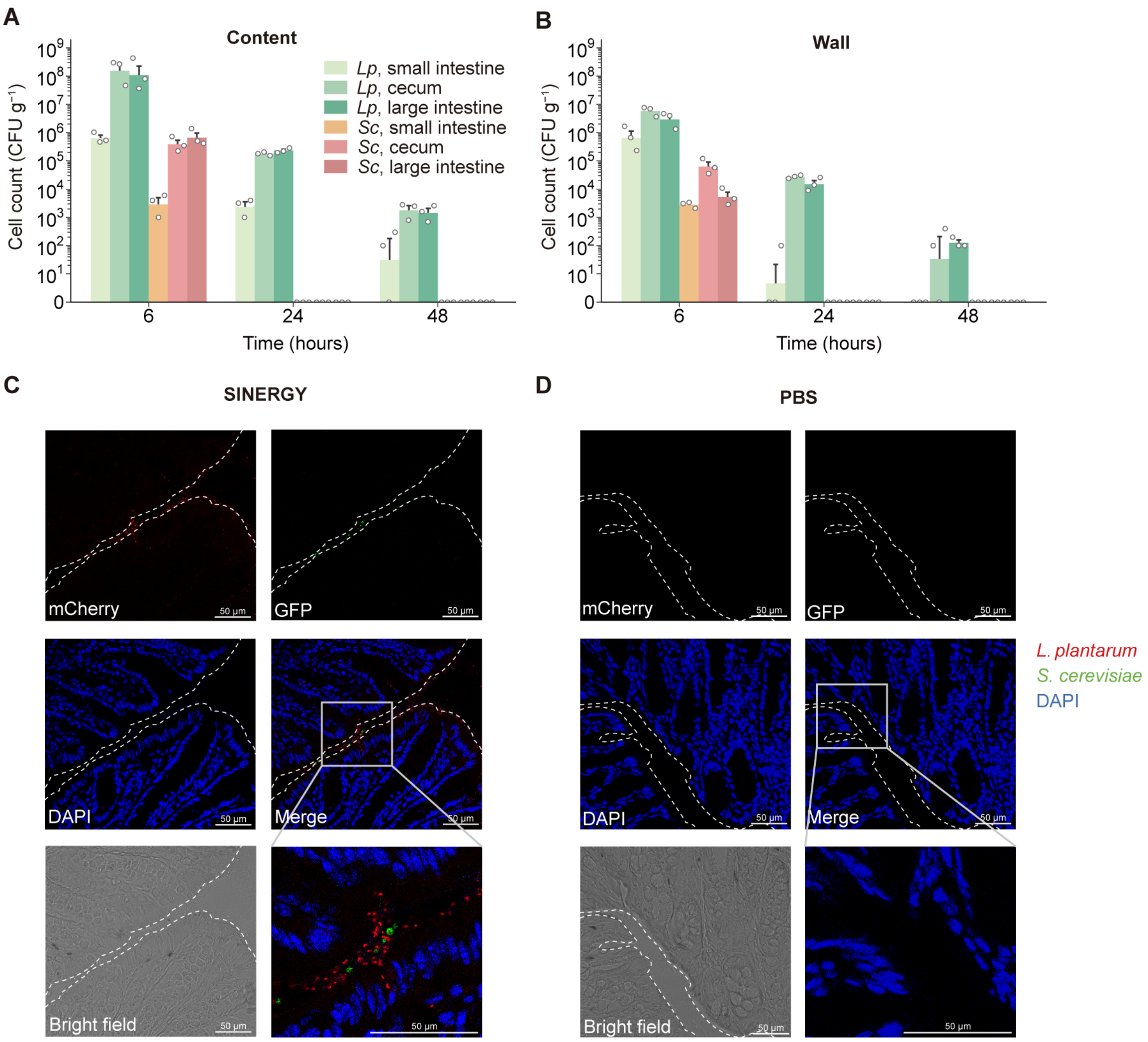


**Fig. S7: Viability of *L. plantarum* and *S. cerevisiae* delivered by SINERGY in the murine gut.**

(**A** and **B**) Microbial cell counts in gut contents (**A**) and on walls (**B**) (mean ± SEM, n = 3 mice). (**C** and **D**) Additional confocal visualization of fluorescent microorganisms in the colon of mice administered with SINERGY (**C**) compared with PBS (**D**).

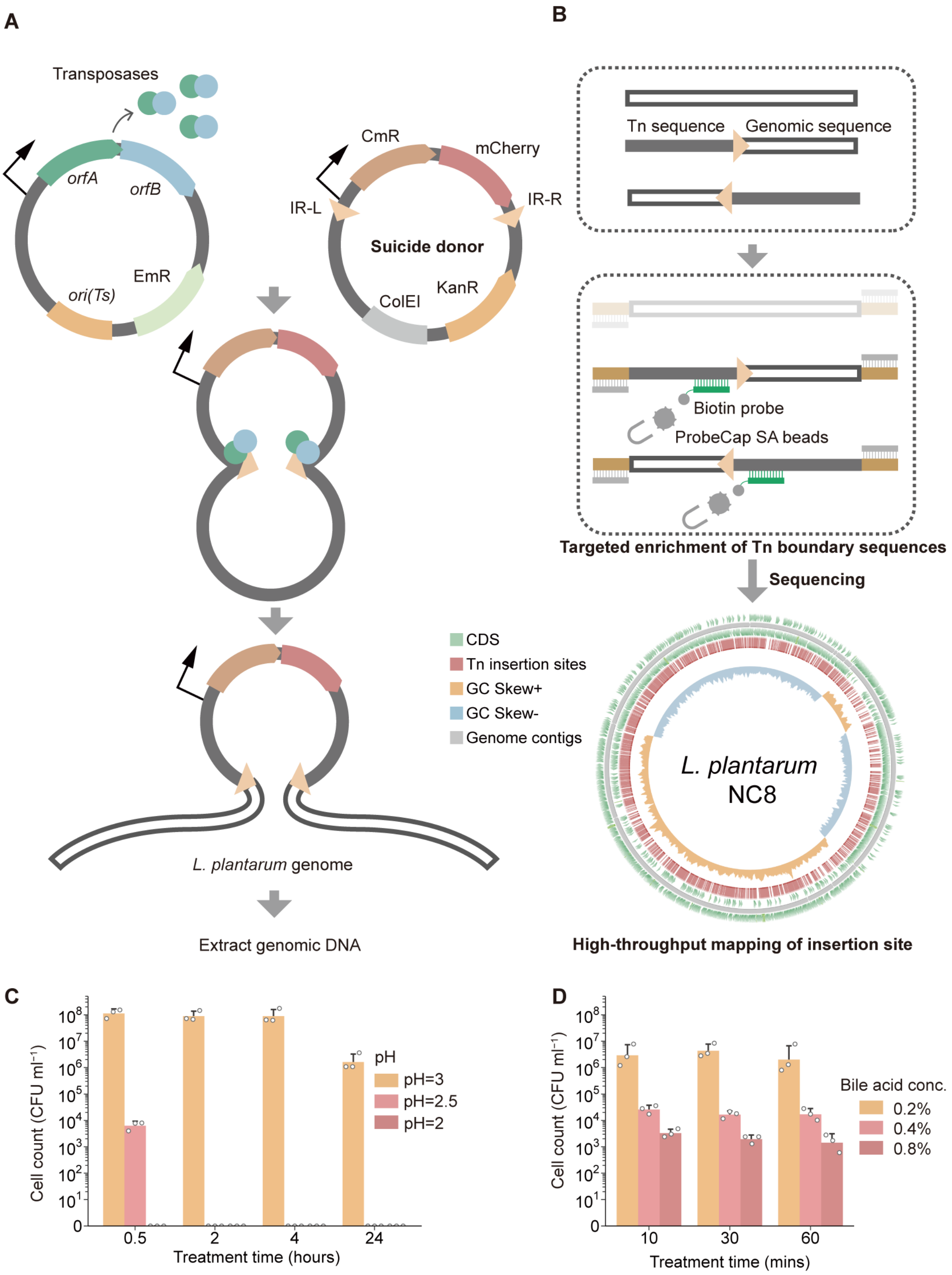


**Fig. S8: Construction of *L. plantarum* Tn-mutagenesis system.**
(**A**) Schematic of the Tn-mutagenesis system in *L. plantarum*. A thermo-sensitive plasmid for constitutively expressing transposases is first introduced into *L. plantarum*, followed by electroporation of a suicide transposon donor that cannot replicate in *L. plantarum* (**Materials and methods**). The integrated colonies that maintain CmR (chloramphenicol resistance) are selected. (**B**) Targeted enrichment sequencing for identifying the Tn-inserted genomic features. (**C** and **D**) Preliminary exploration of screening conditions for low pH treatment (**C**) and bile acid concentration (**D**) (mean ± sd, n = 3 independent experiments).

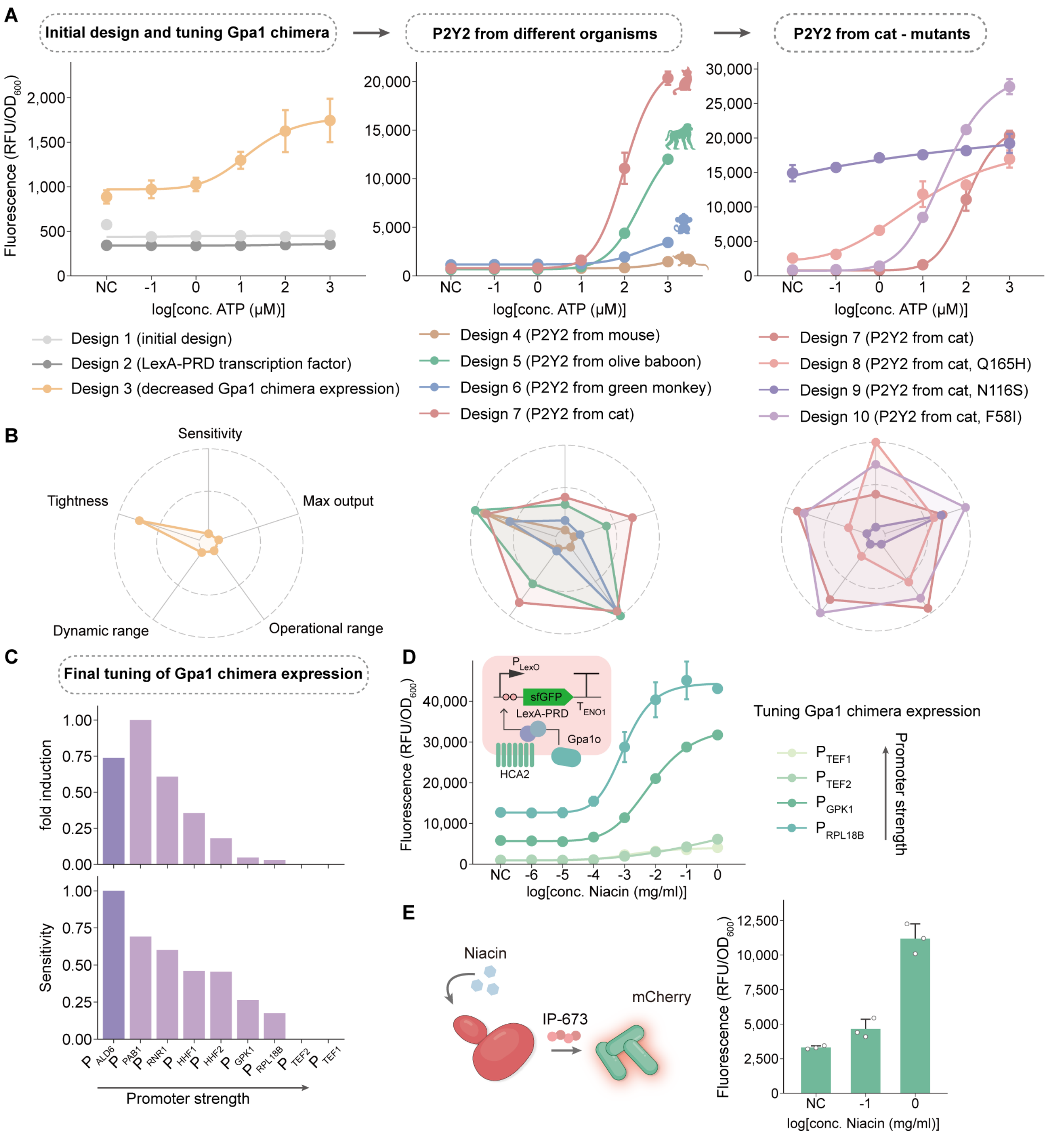


**Fig. S9: Construction and optimization of yeast GPCR-based sensor.**

(**A** and **B**) Dose-response curve (**A**) and radar chart (**B**) of the yeast eATP sensor constructs (mean ± sd, n = 2 or 3 independent cultures).

(**C**) Characterization of eATP sensor fold induction and sensitivity with various promoters for Gα chimera expression (mean).

(**D**) Dose-response curve of yeast GPCR-based niacin sensor designs (mean ± sd, n = 3 independent cultures).

(**E**) Characterization of niacin sensing consortia with *L. plantarum* expressing mCherry as output (mean ± sd, n = 3 independent cultures). NC, negative control.

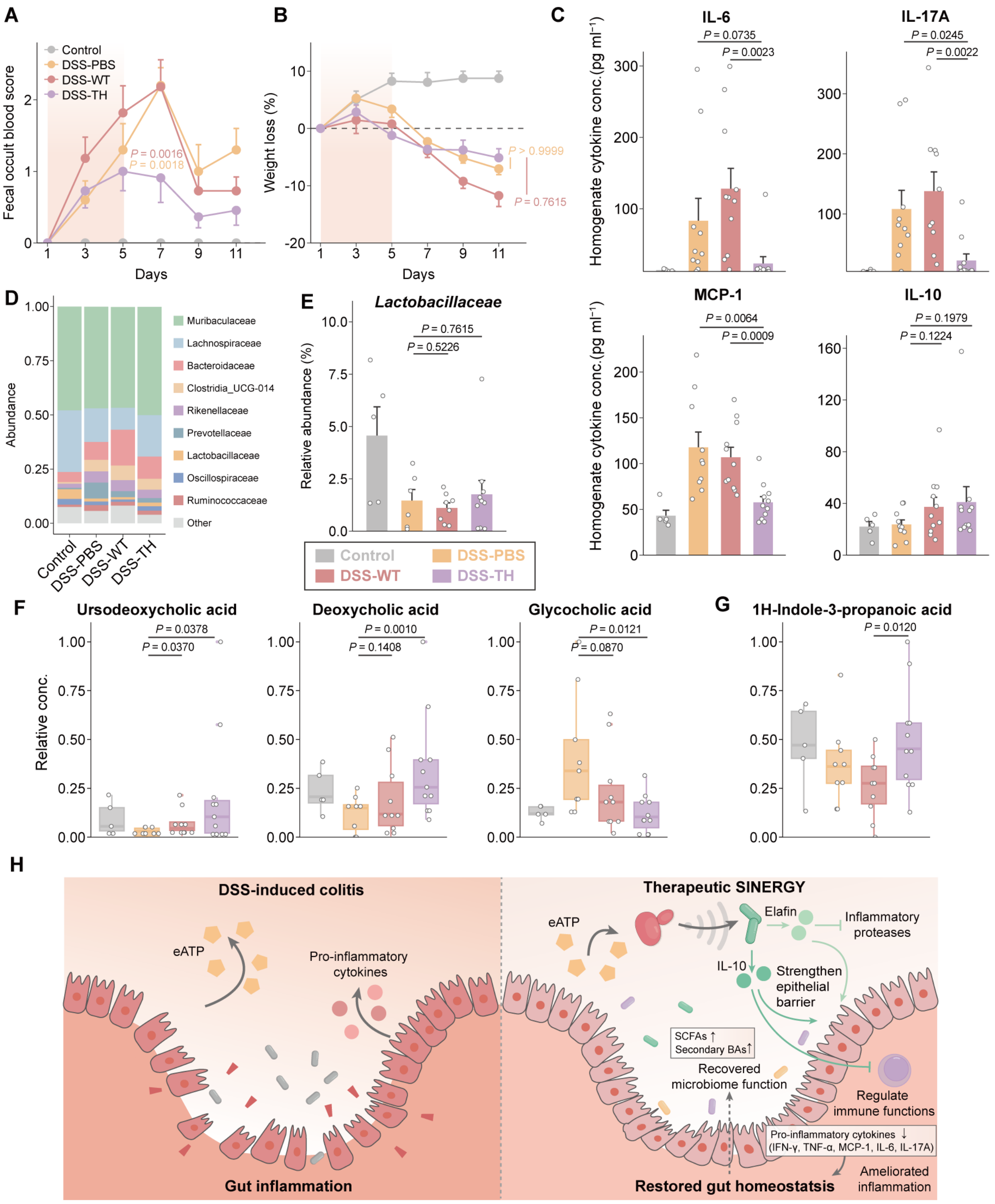


**Fig. S10: Response of DSS-induced colitis to therapeutic SINERGY, consisting of engineered consortia, as revealed by restored microbiota and metabolite levels.**
(**A** and **B**) Fecal occult blood score (**A**) and body weight loss (**B**) of each group of mice (mean ± SEM, statistical significance was determined by two-way ANOVA, P value representing comparison of DSS-TH group versus DSS-PBS or DSS-WT group, n = 5 mice for Control group, n = 10 for DSS-PBS group, n = 11 for DSS-WT and DSS-TH group).
(**C**) Additional cytokine levels of colon homogenate (mean ± SEM, statistical significance was determined by unpaired two-sided t-test, n = 5 samples for Control group, n = 10 for DSS-PBS group, n = 11 for DSS-WT and DSS-TH group).

(**D**) Relative abundance of bacteria classified at a family-level taxonomy.

(**E**) Relative abundance of the *Lactobacillaceae* family (mean ± SEM, statistical significance was determined by unpaired two-sided t-test, n = 5 samples for Control group, n = 7 for DSS-PBS group, n = 8 for DSS-WT and n = 10 DSS-TH group).

(**F** and **G**) Additional bile acid metabolites (**F**) and indole-3-propanoic acid (**G**) of colon content (statistical significance was determined by unpaired two-sided t-test, n = 5 samples for Control group, n = 9 for DSS-PBS group, n = 10 for DSS-WT and n = 11 DSS-TH group).

(**H**) Schematic of the multiple functions of therapeutic SINERGY in DSS-induced colitis.
